# Dynamic rotation of the protruding domain enhances the infectivity of norovirus

**DOI:** 10.1101/2019.12.16.878785

**Authors:** Chihong Song, Reiko Takai-Todaka, Motohiro Miki, Kei Haga, Akira Fujimoto, Ryoka Ishiyama, Kazuki Oikawa, Masaru Yokoyama, Naoyuki Miyazaki, Kenji Iwasaki, Kosuke Murakami, Kazuhiko Katayama, Kazuyoshi Murata

## Abstract

Norovirus is the major cause of epidemic nonbacterial gastroenteritis worldwide. Lack of structural information on infection and replication mechanisms hampers the development of effective vaccines and remedies. Here, using cryo-electron microscopy, we show that the capsid structure of murine noroviruses changes in response to aqueous conditions. By twisting the flexible hinge connecting two domains, the protruding (P) domain reversibly rises off the shell (S) domain in solutions of higher pH, but rests on the S domain in solutions of lower pH. Metal ions help to stabilize the latter conformation in this process. Furthermore, in the resting conformation, the cellular receptor CD300lf is readily accessible, and thus infection efficiency is significantly enhanced. Two P domain conformations were also found in the human norovirus GII.3 capsid. These results provide new insights into the infection mechanisms of the non-envelope viruses that function in dramatic environmental changes such as the digestive tract.

**Significance Statement:** The capsid structures of caliciviruses have been reported to be classified into two different types, according to species and genotype. One is the rising P domain type as shown in human norovirus GII.10 and rabbit hemorrhagic disease virus, where the P domain rises from the S domain surface. The other is the resting P domain type as shown in human norovirus GI.1, sapovirus and San Miguel sea lion virus, where the P domain rests upon the S domain. Here, we demonstrate that the P domain of the murine norovirus infectious particles changes reversibly between the rising and resting P domain types in response to aqueous conditions. Our findings provide new insights into the mechanisms of viral infection of caliciviruses.

## Introduction

Human norovirus (HNV) is a major cause of epidemic nonbacterial gastroenteritis worldwide (1). HNV often causes severe sporadic infections, especially in nurseries and nursing homes. However, there are no efficient treatments or vaccines due to lack of a robust cultivation system for studying the virus (2). Murine norovirus (MNV), a species of norovirus affecting mice, was identified in 2003 (3). MNV propagates in cell lines (3, 4), shares genetic features with, and has biochemically similar properties to HNV (5). For these reasons, MNV has been widely used as a model for HNV infection.

Norovirus is a non-enveloped, positive-stranded RNA virus belonging to the *Caliciviridae* family (6). The common capsid structure of 180 copies of VP1 capsid protein with T=3 icosahedral symmetry is shared in all caliciviruses and has been reported in HNV virus-like particles (VLPs) produced by a baculovirus expression system (7). To form this structure, VP1 is placed in three quasi-equivalent positions of asymmetric units designated as A, B, and C monomers. VP1’s A and B dimers (A/B dimers) are located around the icosahedral fivefold axes, and VP1’s C and C dimers (C/C dimers) are on the icosahedral twofold axes (8). The VP1 protein consists of a shell (S) and protruding (P) domain. The S domain, containing an eight-stranded β sandwich structure, forms a contiguous icosahedral backbone to protect the central viral genome and has high amino acid sequence homology within the *Caliciviridae* family (9). On the other hand, the P domain extended from the S domain via a flexible hinge loop further divided into a lower P1 subdomain and an upper P2 subdomain. With respect to the amino acid sequence, the P1 subdomain can be divided into two parts, the N-terminal P1 part (P1-1) and the C-terminal P1 part (P1-2), with the P2 subdomain interposed. The P2 subdomain is exposed on the top of the P domain and functions for virus attachment to cells (10).

The P domain in caliciviruses shows two conformations (11). The first, called here the rising conformation, is shown in MNV-1 (12), HNV GII.10 (13), and rabbit hemorrhagic disease virus (RHDV) (14, 15), where the P domain rises from the S domain surface, forming an outer shell. The second, called here the resting conformation, is shown in HNV GI.1 (7), sapovirus (9), San Miguel sea lion virus (SMSV) (16) and feline calicivirus (FCV) (17, 18), where the P domain rests upon the S domain. The potential for dynamic structural changes in capsids has been discussed for viral infection and replication, but no direct evidence has been found thus far (11).

Here, using cryo-electron microscopy (cryo-EM), we show that the P domain of MNV infectious particles reversibly rotates ∼70°, in response to aqueous conditions, taking two different conformations; the rising and resting P domain conformations. The P domain extends away from the S domain surface in solutions with higher pH, and rests on the S domain surface in solutions with lower pH. Metal ions help to stabilize the resting conformation in this process. Furthermore, significant differences were found in the two P domain conformations with respect to MNV infection of cultured cells. High-resolution cryo-EM structural analysis using MNV-VLP revealed the structural similarity between infectious particle and VLP in MNVs and elucidates the molecular mechanism of P domain rotation. Our findings provide new insights into the mechanisms of viral infection of the non-enveloped viruses.

## Results

### Dynamic rotation of the protruding domain in MNV controls viral infection

For our structural studies, infectious particles of MNV type 1 (MNV-1) were produced by a reverse genetics system (19), propagated in RAW264.7 cells, and stored in DMEM (Dulbecco’s Modified Eagle Medium). Viral particles were subjected to analysis by single-particle cryo-EM using a 200kV transmission electron microscope (TEM), and a 3D model was reconstructed at 5.3 Å resolution, in which the P domain was stabilized with an outer P2 subdomain-based interaction and rested on the S-domain (Fig. 1*A* and *C* and SI Appendix, Fig. S1*A-D*).

**Fig. 1.**
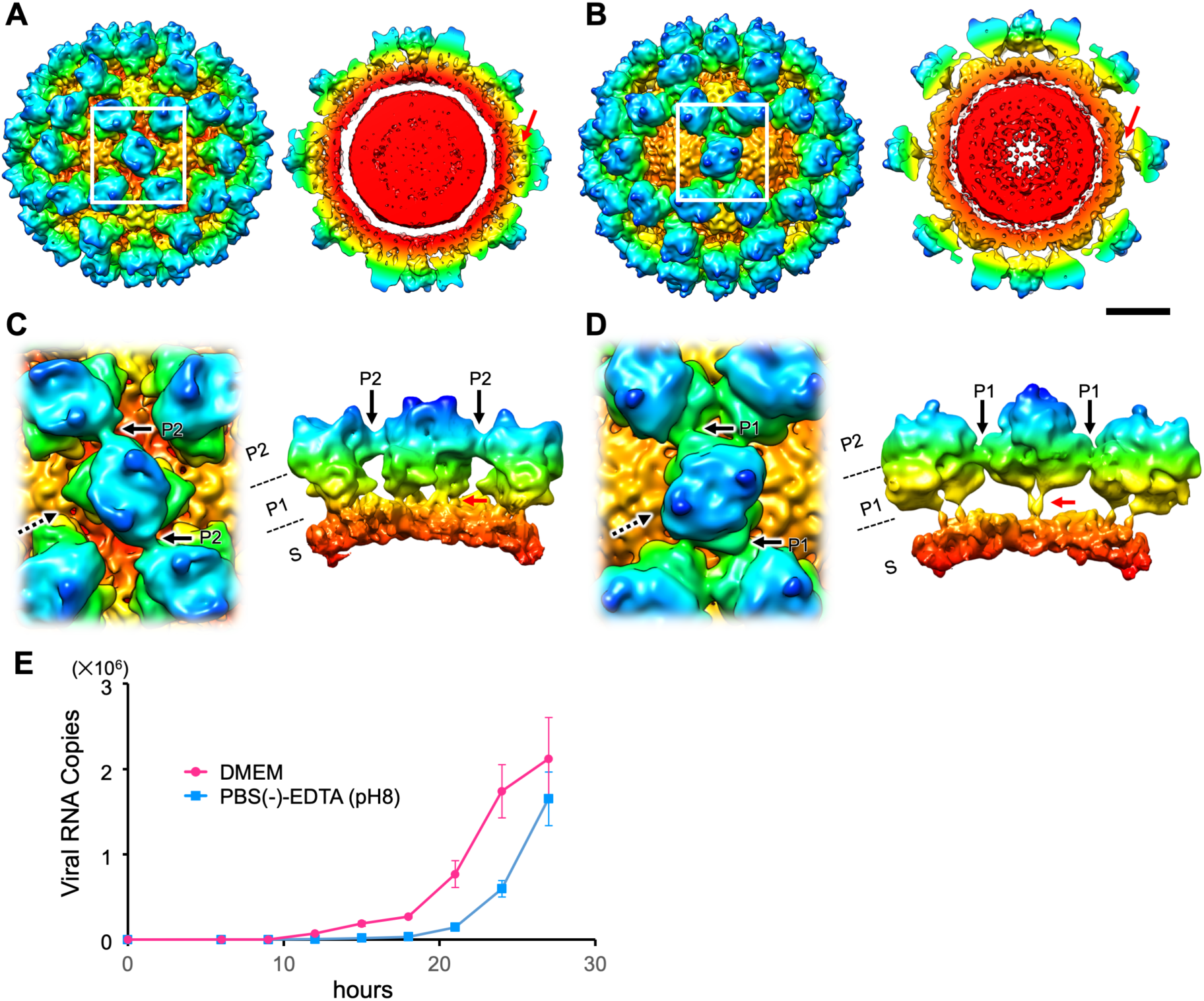
Dynamic rotation of the P domain in MNV-1 infectious particles controls viral infection. (*A* and *B*) Cryo-EM structures of the MNV-1 infectious particles suspended in DMEM (*A*) and PBS(-)-EDTA (pH 8). The maps are low-pass filtered to 8 Å resolution to highlight the domain structure. The left and right panels are the isosurface representation and the center section, respectively. (*C* and *D*) Left panels are the enlarged views of the boxes in *A* and *B*. Right panels are views from the direction of the dotted arrows shown in the left panel. The P domain of MNV-1 suspended in DMEM rests on the S domain (red arrows in *A* and *C*) and interacts with the adjacent P domains at the outer P2 subdomain level (black arrows in *C*), called the resting P domain conformation. By contrast, the P domain of MNV-1 suspended in PBS(-)-EDTA (pH 8) rises off the S domain (red arrows in *B* and *D*) and interacts with the adjacent P domains at the inner P1 subdomain level (black arrows in *D*), called the rising P domain conformation. In the rising P domain conformation, all P domain dimers rotate ∼70° clockwise, compared to the resting P domain conformation, and the interaction sites are changed from the P2 to the P1 level. The P domain rising off from the S domain in PBS(-)-EDTA (pH 8) reversely rests on the S domain in DMEM. Scale bar, 100 Å. (*E*) Propagation curves of MNV-1 infectious particles pretreated with DMEM and PBS(-)-EDTA (pH 8), respectively. MNV-1 infectious particles were used to infect RAW264.7 cells and the number of the duplicated viral RNA copies along the timeline were measured by qRT-PCR. Error bars represent the standard deviations. MNV-1 infectious particles pretreated with PBS(-)-EDTA (pH 8) propagated slower than those pretreated with DMEM.

Our results differed from a previous report of the 8 Å cryo-EM structure of MNV-1 suspended in PBS (14), in which the P domains rotated ∼70° clockwise were mutually stabilized by interactions based on the inner P1 subdomain, and extended away from the S domain (14). To investigate these structural inconsistencies, we suspended the infectious particles in various aqueous solutions and examined them by cryo-EM. We were able to reproduce the reported MNV-1 capsid structure at 7.3 Å resolution when the viral particles were suspended in a PBS(-) solution containing 20m M EDTA at pH 8.0 (PBS(-)-EDTA (pH 8.0)) (Fig. 1*B* and *D*, SI Appendix, Fig. S1*E-H*). We found that the previously reported structure (14) of MNV-1 appears when the pH was above pH 7 and metal ions were removed from solution by chelation with EDTA (SI Appendix, Fig. S1*I*). Furthermore, the dynamic rotation of the P domain of MNV-1 infectious particles occurred reversibly in response to aqueous conditions.

Next, we sought to elucidate the physiological functions of the two P domain conformations in terms of infectivity of the viral particles. When MNV-1 infectious particles pretreated with PBS(-)-EDTA (pH 8) were infected to RAW264.7 cells cultured in DMEM (pH 7.2 −7.4) (see Materials and Methods), they showed significantly more propagation delay (∼3 hours) than those pretreated with DMEM (Fig. 1*E*). To investigate the reason for the propagation delay, we first compared the virus attachment on the host cell surface at 30 minutes after infection using quantitative real-time RT-PCR (qRT-PCR). The binding of MNV-1 particles pretreated with PBS(-)-EDTA (pH 8) to RAW264.7 cells did not show a signxificant difference from those pretreated with DMEM (Left bars in SI Appendix, Fig. S2*A*). RAW264.7 cells are known to express several molecules involved in MNV adsorption (20). In contrast, the HEK293T cells in which CD300lf is genetically expressed (HEK293T/CD300lf) can adsorb MNV particles on the cell surface through direct interaction with the expressed CD300lf (21). Therefore, the HEK293T/CD300lf cells were also used to evaluate MNV adsorption depending on the two P domain conformations. On the HEK293T/CD300lf cells, MNV-1 particles pretreated with PBS(-)-EDTA (pH 8) significantly showed a reduction of the cell binding compared to the particles pretreated with DMEM (Right bars in SI Appendix, Fig. S2*A*). In addition, flow cytometry (fluorescence-activated cell sorting (FACS)) analysis demonstrated that MNV-1 particles pretreated with PBS(-)-EDTA (pH 8) showed lower binding on the cells (Median=268 CV=157) compared with the MNV-1 particles pretreated with DMEM (Median= 357, CV=275) when the virus particles mixed with the cells at MOI (Multiplicity of infection) = 10 (SI Appendix, Fig. S2*B*). Furthermore, we compared early genome replication in RAW264.7 cells from 30 minutes to 12 hours. The virus genome of MNV-1 pretreated with PBS(-)-EDTA (pH 8) increased only 3.6 fold, while the virus genome of MNV-1 pretreated with DMEM increased 26.4 fold (SI Appendix, Fig. S2*C*).

We also identified using cryo-EM that the rising P domain conformation gradually changed to the resting conformation in 2 to 8 hours, when MNV-1 particles pretreated with PBS (-)-EDTA (pH 8) were suspended in DMEM (pH 7.2 – 7.4) (SI Appendix, Fig. S3*A*). Interestingly, particles mixed with two conformations were also observed between 2 and 6 hours (SI Appendix, Fig. S3*B*). The mixed P domain structure within a single particle suggests that even if the conversion of individual P domains may be rapid, it would take more time to modify the overall conformation of the capsid, where the P domains interact and are connected together like a net. Consequently, the delay time of the virus propagation pretreated with PBS(-)-EDTA (pH 8) possibly corresponds to the time of the overall P domain conformational change of the virus particles in the cell culture media. These results suggest that the rising conformation of the P domain prevents the initial viral binding to the host cell surface via CD300lf, and it takes time to retrieve infectivity by changing the rising conformation to the resting conformation in the whole capsid. Hence, MNV-1 can control viral infectivity by the dynamic rotation of the P domain.

The same dynamic rotation of the P domain was also observed with MNV type S7 (MNV-S7) (SI Appendix, Fig. S4 and S5). MNV-S7 is a norovirus isolated from mouse stools in Japan in 2007 (22) and shows similar pathogenicity to MNV-1. The amino acid sequence of MNV-S7 VP1 protein shows 97% homology with MNV-1, with the nonhomologous 18 amino acids concentrated in the P domain (SI Appendix, Fig. S6). Interestingly, in the case of MNV-S7, rotation and elevation of the P domain only occurred in the solution of PBS(-)-EDTA at pH 8 or higher (SI Appendix, Fig. S4*K*), which is slightly higher than that of MNV-1 (pH 7 or higher (SI Appendix, Fig. S1*I*)). Viral propagation was also delayed for viruses pretreated with PBS(-)-EDTA (pH 8) solution compared to viruses pretreated with DMEM, but the difference was smaller than that of MNV-1 (Fig. 1*E* and SI Appendix, Fig. S4*L*). The results indicate that the MNV-S7 particles more readily convert conformation to the resting state than the MNV-1 particles in the infection medium of DMEM (pH 7.2 – 7.4). However, the binding of MNV-S7 particles pretreated with PBS(-)-EDTA (pH 8) to both RAW264.7 and HEK293T/CD300lf cells showed a significant difference from those pretreated with DMEM (SI Appendix, Fig. S5*A*). For early genome replication in RAW264.7 cells from 30 minutes to 12 hours, the virus genome of MNV-S7 pretreated with PBS(-)-EDTA (pH 8) increased only 7.4 fold, while the virus genome of MNV-S7 pretreated with DMEM increased 17.2 fold (SI Appendix, Fig. S5*B*). The binding mechanism of MNV-S7 to host cells may be slightly different from that of MNV-1.

### Capsid structure of MNV-S7 VLP

To investigate the molecular mechanism of the dynamic rotation of the P domain in the MNV capsid, we determined the atomic structure of the entire MNV capsid by single-particle cryo-EM in a 300kV TEM. VLPs of MNVs were used for this structural analysis. The higher resolution capsid models are easier to obtain from VLPs than infectious particles containing nucleotides (23), and VLPs satisfy the biosafety level requirements of the 300kV EM room. VLPs of MNV-1 and MNV-S7 were prepared using a modified baculovirus expression system (24, 25) and stored in DMEM. As a result, VLPs of MNV-1 formed multiple types of icosahedral particles of 40–50 nm, whereas those of MNV-S7 formed a uniform icosahedral particle in the size of about 40 nm. Therefore, VLPs of MNV-S7 were used for the high resolution cryo-EM analysis, and the 3D model was determined at 3.5 Å resolution (Fig. 2 and SI Appendix, Fig. S7). Local resolutions were in the range of 3.3 to 3.7 Å in the capsid (Fig. 2*B*). The highest resolution of 3.3 Å was observed in the S domain, and lower resolutions were mainly located in the peripheral regions of the P domain.

**Fig. 2.**
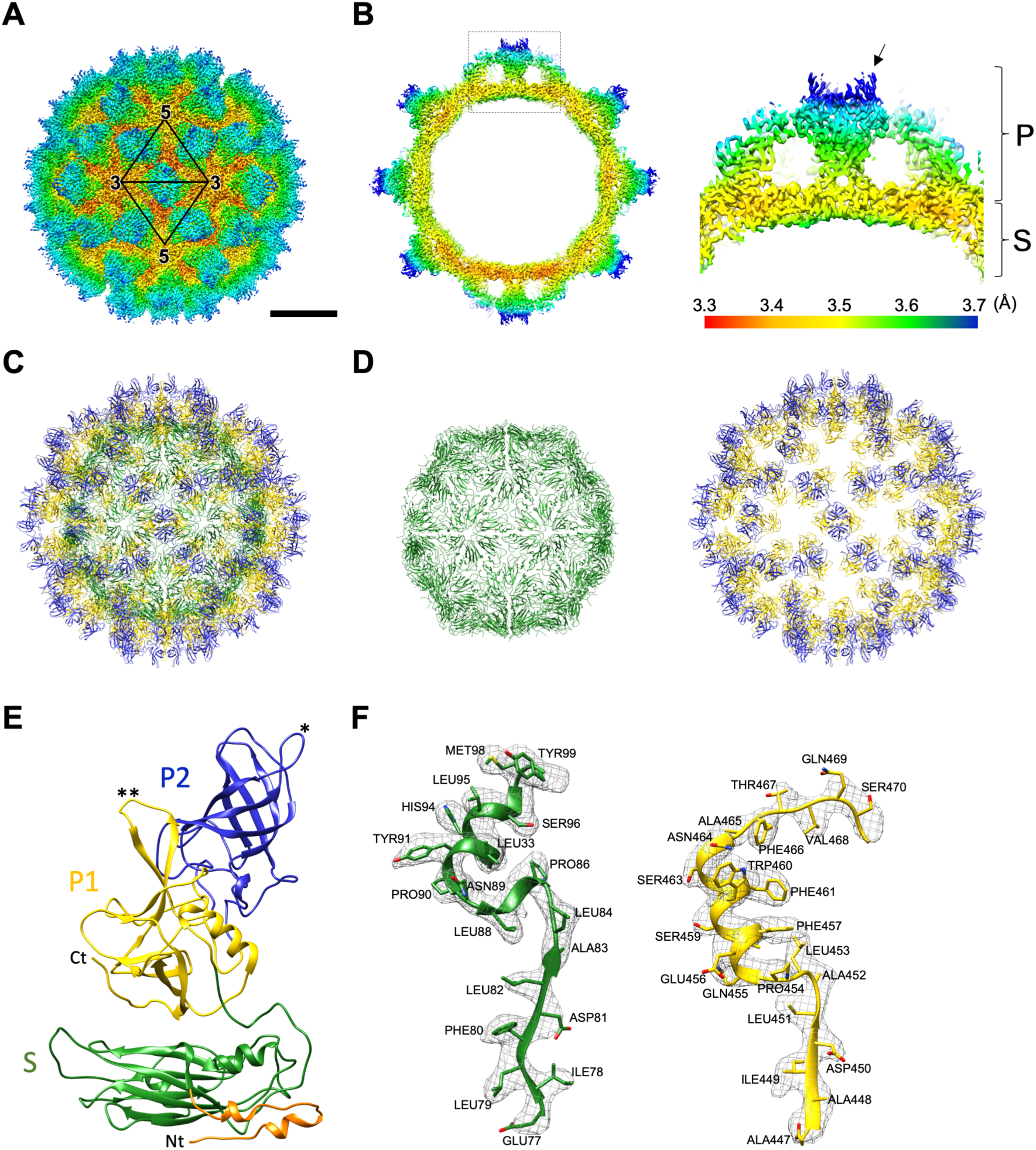
Cryo-EM structure of MNV-S7 VLP at 3.5 Å resolution. (*A*) Cryo-EM map of MNV-S7 VLP in DMEM viewed down the icosahedral twofold axis. Coloring is based on radii, as follows: red, up to 161 Å; yellow 161–171 Å; green, 171–191 Å; cyan, 191–211 Å; blue, 211 Å and above. Scale bar, 100 Å. (*B*) Local resolution assessment on a cross section of the MNV-S7 VLP. The right panel shows a higher magnification of the square in the left image; the P and S domains are labeled. Coloring is based on the local resolution at the bottom. Arrow indicates a horn-like structure in the P domain. (*C* and *D*) The Cα backbone of the S+P (*C*), S (the left panel in *D*), and P (the right panel in *D*) domains in the icosahedral MNV-S7 particle at the same orientation with *A*. The coloring follows the standard designation of P1 subdomain (yellow), P2 subdomain (blue) and S domain (green). (*E*) A ribbon model of the *C* monomer of VP1 in the MNV-S7 capsid. 502 amino acids from Val30 to Gly531were modeled in the C monomer, including the P1 subdomain (yellow), P2 subdomain (blue) and S domain (green). Single and double asterisks show the loops, which function to stabilize the P domain dimer. (*F*) Representative cryo-EM electron densities of several amino acids and the fitted atomic models.

Atomic models of the MNV-S7 VLP capsid were built for the quasi-equivalent A, B, and C monomers of VP1, respectively (SI Appendix, Fig. S6 and Table S2). The complete Cα backbone structure of the MNV-S7 VLP is shown in Fig. 2*C*. Other features of the models (e.g., S and P domains, Cα backbone of the VP1 C monomer) are presented in Fig. 2*D-F*. For modeling VP1’s A and B monomers, 513 amino acids from Gln19 to Gly531 and 516 amino acids from Ala16 to Gly531 were used, respectively (SI Appendix, Fig. S6). For modeling VP1’s C monomers, 502 amino acids from Val30 to Gly531 were used (SI Appendix, Fig. S6). The models were also fitted to the 5.2 and 5.3 Å resolution cryo-EM maps of the MNV-S7 and MNV-1 infectious particles with high cross-correlation coefficients of 0.92 and 0.91 (see Materials and Methods), respectively, indicating that the structure of MNV-S7 VLP is highly similar to both MNV-S7 and MNV-1 infectious particles.

A comparison of the atomic model of the MNV-S7 VLP with the reported crystallographic models of the MNV-1 P domain (PDB ID: 3LQ6 and 3LQE) showed a high degree of structural similarity, except for two flexible loops. The βC’’–βD’’ loop (SI Appendix, Fig. S6) extends outward and forms a part of the horn-like structure (22) (arrow in Fig. 2*B*) on the top of the P domain in our cryo-EM model (single asterisk in Fig. 2*E*), whereas the crystallographic models of MNV-1 extends horizontally with respect to the capsid surface (26). Another loop located on the opposite side of the βC’’–βD’’ loop conversely extends horizontally with respect to the capsid surface in our cryo-EM model (double asterisk in Fig. 2*E*), whereas that of the crystallographic model extends outward (26). These two loops were disordered in the recent reported crystallographic model of the MNV-1 P domain dimer containing the soluble domain of the cellular receptor, CD300lf (PDB ID: 5OR7) (27). In our cryo-EM model, these loops interact with each other in the VP1 dimer and function to stabilize the P domain dimers in the viral capsid.

### Molecular interactions between VP1 proteins

To understand the capsid structure of the resting P domain conformation, showing higher levels of infectivity, the molecular interactions between the P domain dimers of the MNV-S7 VLP suspended in DMEM were investigated. The cryo-EM map of the MNV-S7 VLP in DMEM revealed several hydrophobic and charged interactions between the neighboring A/B and C/C dimers at the P2 level (yellow in Fig. 3). We propose the following chemical interactions between residues on the P domain surface: Asn409, Gly411, Leu412, and Pro415 located on the βF’’–βB’ loop (SI Appendix, Fig. S6) of the C/C dimer interact with Gln371, Arg373, Val368, and Pro319 located on the βB’’–βC’’ and βD’’–βE’’ loops (SI Appendix, Fig. S6) of the A/B dimers, respectively (Fig. 3*B* and *C*).

**Fig. 3.**
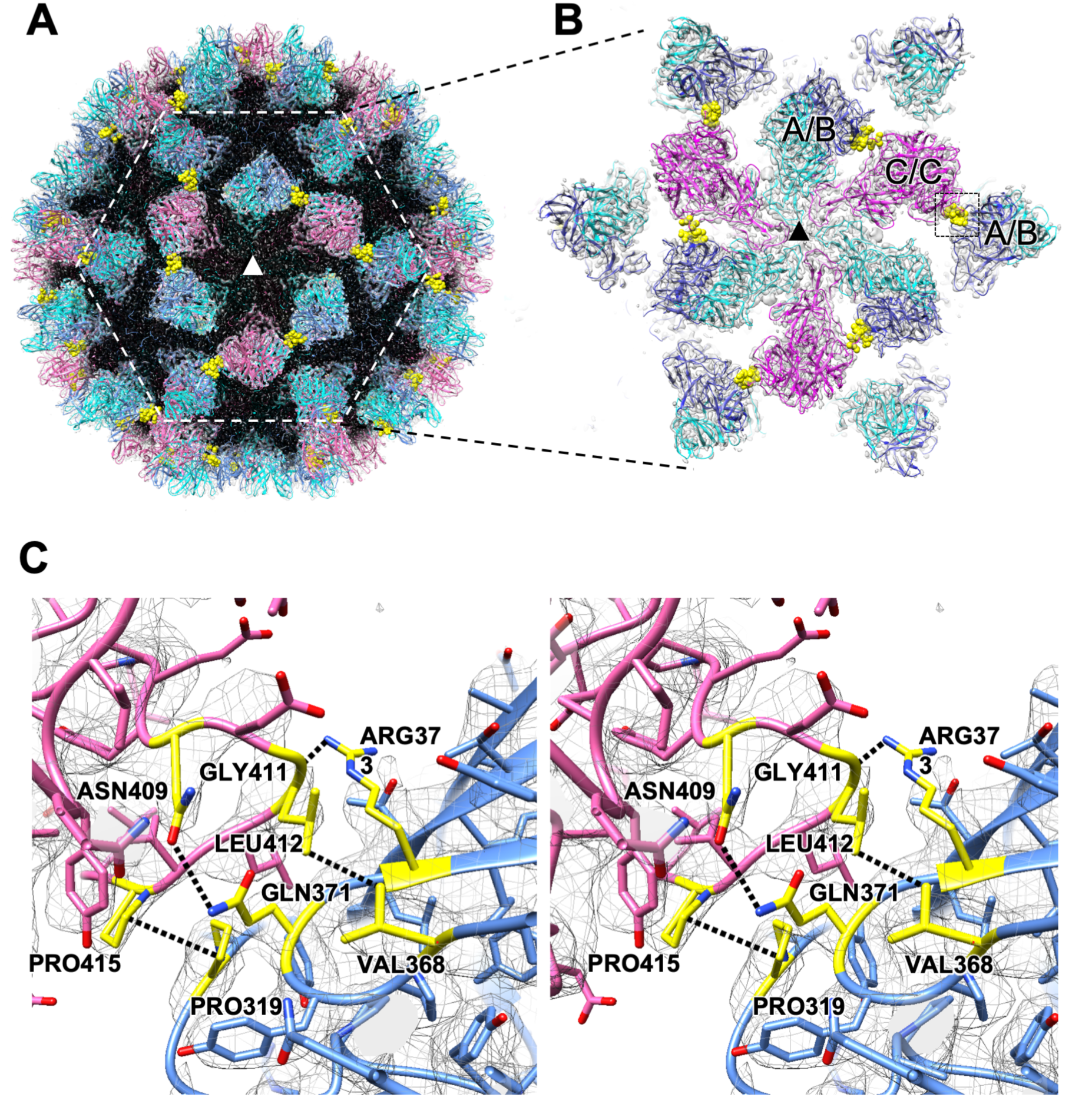
Interactions between the resting P domain dimers. (*A*) A cryo-EM map of the MNV-S7 VLP highlights the interactions of the P domain dimers. The central threefold axis is labeled with a triangle. To emphasize the P-domains, the density map corresponding to the S-domain is labeled in black. (*B*) The enlarged view of the dotted hexagon in *A*. C/C-dimers (purple) on the twofold axis interact with adjacent A/B-dimers (blue) located around the fivefold axis. The residues interacting with the adjacent P domain dimers are colored by yellow. (*C*) A stereo view of the candidate chemical interactions between the C/C and A/B dimers indicated by the dotted square in *B*. Asn409, Gly411, Leu412, and Pro415 of the C/C dimer are facing with Gln371, Arg373, Val368, and Pro319 of the A/B dimer, respectively, where hydrophobic and charged interactions are suggested to be formed.

As expected from sequence homology (SI Appendix, Fig. S6), the structure of the S domain of MNVs was similar to the HNV GI.1 VLP (PDB ID: 1IHM) (7) except for some local distortions (SI Appendix, Fig. S8*G* and *H*). Our cryo-EM structural study showed several elaborate crosslinks between adjacent N-terminals of the S domains, by successfully modeling the residues from Gln19 in the A monomer, Ala16 in the B monomer, and Val30 in the C monomer (SI Appendix, Fig. S8*A*). The flexible N-terminals formed various interactions with each other between adjacent S domains at the twofold, threefold, pseudo-threefold, and fivefold axes (SI Appendix, Fig. S8*B-F*). The basic architecture is similar to the previous reports of other members of the *Caliciviridae* family, however, the fine structure is so far unique. As shown in SI Appendix, Fig S8*G* and *H*, the currently reported N-terminal arms (NTAs) are structurally divided into two groups. One group shown in HNV GI.1 (7) and RHDV (15) is that the NTAs run along the outer bottom edge of their own S domain and interact with the neighboring S domain (Blue, and cyan in SI Appendix, Fig. S8*G* and *H*). The other group shown in SMSV (16) and FCV (17) is that the NTAs run along the outer bottom edge of their neighboring S domain and interact with them (Yellow and green in SI Appendix, Fig. S8*G* and *H*). The NTA in MNV has the similar conformation with the first group (Red in SI Appendix, Fig. S8*G* and *H*). Consequently, the rigid capsid shell of MNV was maintained by these outer P domain interactions and inner S domain crosslinks.

### Molecular mechanism of the reversible rotation of the P domain dimers

We investigated the capsid structure of MNV in the rising P domain conformation to understand the molecular mechanism of the dynamic rotation of the P domain. The atomic model of the MNV-S7 VLP in DMEM was modified and fitted into the 7.2 Å cryo-EM map of the MNV-S7 infectious particle suspended in PBS(-)-EDTA (pH 8) with the high cross-correlation coefficient of 0.92 (SI Appendix, Fig. S4*I*) (see Materials and Methods) and the atomic model of the rising P domain conformation was estimated (SI Appendix, Fig. S9). The new model in PBS(-)-EDTA (pH 8) showed that the P domain rotates clockwise by ∼70° and rises up from the S domain surface by ∼13 Å compared to the original model in DMEM (Fig. 4*A-D*). The cryo-EM map of the rising P domain conformation suggested that new interactions link the P1 subdomain of the C/C dimer to the adjacent P1 subdomain of the A/B dimers via hydrophobic and charged interactions. These are formed between residues on the newly facing P domain surface; i.e., Pro425, Ser504, Leu524, and Gln526 in the βF’’–βB’ and βF’–αG loops, while the C-terminal (SI Appendix, Fig. S6) of the P1 subdomain in the C/C dimer interact with Phe423, Gln263, Leu524, and Arg523 located in the same loops and C-terminal in the A/B dimer, respectively (SI Appendix, Fig. S9*B* and *C*).

**Fig. 4.**
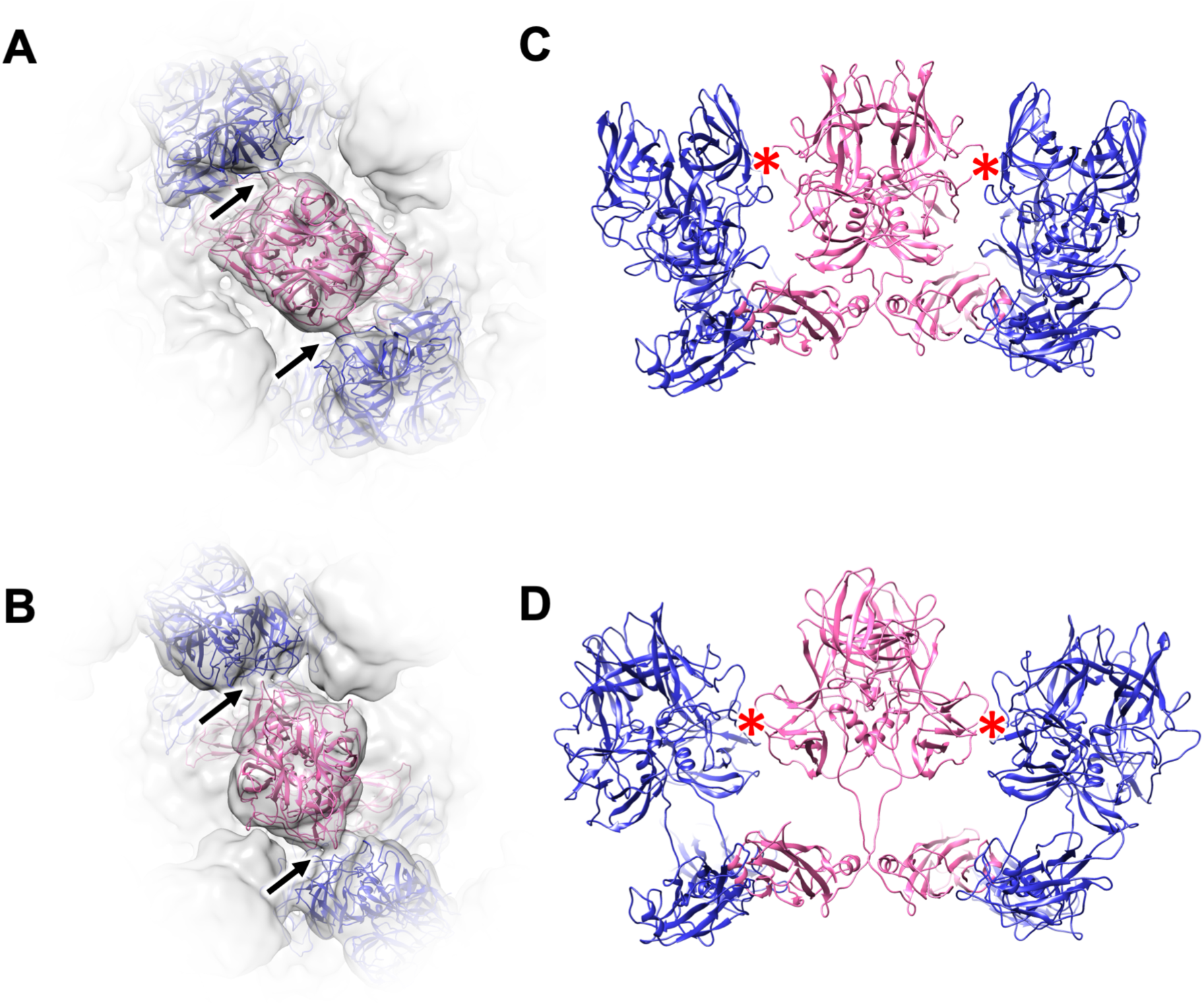
Molecular mechanism of the reversible dynamic rotation of the P domain dimers. (*A* and *B*) Ribbon models of the C/C dimers of MNV-S7, suspended in DMEM and PBS(-)-EDTA (pH 8) are modified and fitted into the map. Arrows indicate where the C/C dimer interacts with the A/B dimers. (*C* and *D*) Side views of the ribbon models showed in *A* and *B*, respectively. The interactions linking the adjacent P domains are indicated by red asterisks. The animated dynamic rotation of the P domain dimers between two conformations are shown in Movie S1.

An animated comparison of the P domain conformations and reversible structural dynamics of the MNV capsid under different aqueous conditions is shown in Movie S1. The P domains in DMEM rested on the S domains and were stabilized by interaction between the adjacent P domains at the P2 level (Fig. 4*A* and *C*), while the P domains in PBS(-)-EDTA (pH 8) rose up from the S domain and were stabilized by interaction between the adjacent P domains at the P1 level (Fig. 4*B* and *D*). The flexibility of the hinge connecting the P and S domains (SI Appendix, Fig. S10*A*) is important for dynamic twisting of the P domain, which moves the P domain up and down by ∼13 Å. In addition, as the P domain rose, the A/B dimer was bent slightly toward the fivefold axis (SI Appendix, Fig. S11).

We also observed two conformations of the P domain in the same HNV GII.3 (TCH04-577 strain) VLPs by cryo-EM (Fig. 5). Single particle analysis showed that the two P domain conformations were similar to the resting and rising P domain conformations of MNVs (Fig. 5, SI Appendix, Fig. S12). In the T=3 icosahedral particles of the HNV GII.3 VLP suspended in DMEM, 16% of the total showed the rising P domain conformation, and the rest showed the resting P domain conformation. The ratio did not change even in PBS(-)-EDTA (pH 8.0). Our observations suggest that the capsid of HNV can also assume two reversible P-domain conformations, though dynamic rotation is currently identified only in MNV infectious particles. Further investigations are needed to determine the factors that control the orientation of the P domain in HNV.

**Fig. 5.**
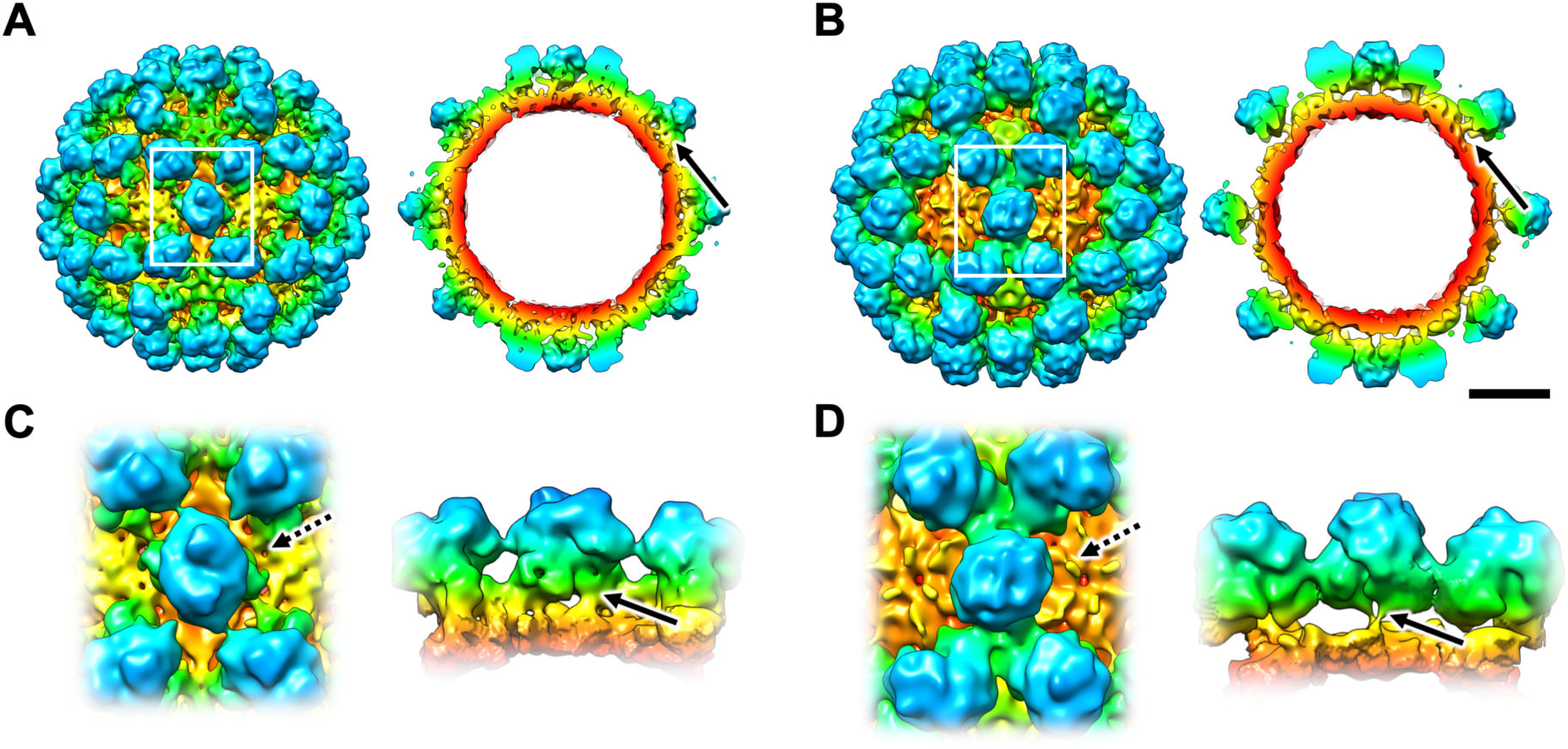
Structures of the resting and rising P domain conformations in HNV GII.3 VLPs. (*A* and *B*) Cryo-EM structures of the HNV GII.3 VLP with the resting and rising P domain conformations (arrows) displayed at 13 Å resolution, respectively. Isosurface and center section images of the whole particles are shown in left and right panels, respectively. Rotation of the P domain by ∼70° and movement of the interaction site result in a conformational change to the HNV P domains. Scale bar, 100 Å. (*C* and *D*) Left panels are enlarged views of the rectangle boxes in *A* and *B*. Right panels are views from the direction of the dotted arrows shown in the left panel. Arrows indicate the resting and rising P domain conformations, respectively.

## Discussion

Here, we reported that reversible dynamic rotation of the P domain results in two conformations of the MNV capsid. The main trigger for the conformational change was the pH of the solution: a higher pH changed the conformation of the P domain to the rising state, while a lower pH changed it to the resting state. Interestingly, the pH thresholds differed by genotype: MNV-1 showed ∼pH 7 (SI Appendix, Fig. S1*I*) and MNV-S7 showed ∼pH 8 (SI Appendix, Fig. S4*K*). However, it was not clear what determines the threshold for interconversion. Some different amino acids between the P and S domains may perform it by changing surface charge, while metal ions binding to the region between the P and S domains, may help to stabilize the resting conformation (SI Appendix, Fig. S10). Thus, in addition to higher pH, EDTA is necessary to remove metal ions from solution, permitting the conversion of the capsid protein to the rising state. As evidence, we identified density between the two carboxylates (Glu223 and Asp231) in the flexible hinge of the resting P domain conformation, where a metal ion is expected to bind to these residues (SI Appendix, Fig. S10*B* and *C*). This suggests that the flexible hinge between the P and S domains could be folded by binding metal ions in DMEM and unfolded by releasing the metal ion in PBS(-)-EDTA (pH 8). In addition, several potential interaction sites connecting the S and P domains are identified in the resting state map (e.g. Glu66 in S domain - Ser519 in P domain, Glu176 in S domain - Tyr227 in P domain, Asp174 in S domain - Gln469 in P domain). These residues may also function to stabilize the resting conformation via metal ions in DMEM. The localization of all metal ions which directly couple the P and S domains will be identified by future higher resolution work.

Our infection and adhesion experiments of the virus particles to the host cells clearly suggested that the viruses in the resting P domain conformation can attach to and infect the host cells, but the viruses in the rising P domain conformation cannot. Further, we performed cryo-EM single particle analysis of MNV-S7 VLP incubated with recombinant CD300lf (rCD300lf) molecules, which is the soluble domain of a cellular receptor of MNV discovered recently (21, 28). The VLPs were treated in DMEM or PBS(-)-EDTA (pH 8) to induce the resting or rising P domain conformation before incubation with rCD300lf. The cryo-EM map of the MNV-S7 VLP pretreated in DMEM showed a weak density on the side of the A/B dimer and extending towards the center of the threefold axis in the icosahedral capsid, while the map of the MNV-S7 VLP pretreated in PBS(-)-EDTA (pH8) was not. Because of the weak density where rCD300lf did not appear to bind to all P-domains of the icosahedral particles, the map was further processed with the “particle symmetry expansion” technique (see Materials and Methods). The individual P-domain dimers on the capsid were classified in 3D, and the only 3D class with the density assumed to be bound by rCD300lf (5.6% of the total) was selected (SI Appendix, Fig. S13*A*). The selected P-domain dimers were then used for 3D refinement by imposing C1 symmetry. As a result, the extra density on the P domain dimer became clearer, though the overall density of rCD300lf has not yet been acquired (SI Appendix, Fig. S13*B-C*).

In order to investigate the binding mode of rCD300lf to the P domain, a protein-protein docking simulation was performed using the MOE program (see Materials and Methods) (SI Appendix, Fig. S13*D*-*E*). In the docking simulation, and based on the extra density that appeared upon addition of rCD300lf (purple in SI Appendix, Fig. S13*C*), Ser43 and Lys45 in the C-C’ loop of rCD300lf interacted with Gly400 in the βF’’-βB’ loop and Ala446 in the βB’-βC’’ loop (SI Appendix, Fig. S6) of the MNV-S7 P domain, respectively, through hydrogen bonding (SI Appendix, Fig. S13*E-F*). The adaptation of our rCD300lf binding model in our cryo-EM map showed that the accessibility of CD300lf to the P domain is altered in the resting and rising P domain conformations. The approach of rCD300lf to the P domain is readily accessible in the resting P domain conformation (SI Appendix, Fig. S13*G*), whereas it is spatially restricted in the rising P domain conformation (SI Appendix, Fig. S13*H* and Movie S2). Although this poor accessibility is unlikely to be a direct cause of non-infectivity of the rising P domain conformation, it showed a possibility that the rotation of the P domain modulates viral infectivity by altering the accessibility of the viral epitopes to cellular receptors.

The binding mode of rCD300lf on the P domain were slightly different from the results of the recent X-ray crystallography using the co-crystal (PDB ID: 5OR7, 6C6Q) (27, 29) (SI Appendix, Fig. S13*I*). In the model, the actual binding site was close, but rCD300lf itself approached from a higher position with a rotation of ∼90°. The rCD300lf engaged a cleft between the A-B and D-E loops of the P2 subdomain in the crystal model (29), while the position in our model is slightly below the two loops. This difference was likely due to the different binding methods or the limited resolution of our cryo-EM map. However, although the binding mode is different, the spatial restriction of the CD300lf accessibility depending on the dynamic P domain rotation can be applied to the co-crystal model.

The resting P domain conformation showed higher accessibility of the cellular receptor CD300lf than the rising P domain conformation, causing greater viral infectivity. These observations raise a question: why does norovirus require such a reversible rising P domain conformation? One possibility is that the conformational changes are important during virus assembly and disassembly in host cells. Tobacco mosaic virus (TMV), well-studied for its structural and physiological characteristics, forms a metastable structure in the cell, where lower concentration of metal ions and higher pH are proposed as the trigger for virus disassembly (30). A recent cryo-EM study showed the calcium ion-dependent conformational changes of TMV, indicating that chemical bonds involving calcium ions prevents disassembly of TMV (31). Dengue and Zika viruses also show pH-dependent structural changes associated with viral replication (32, 33). In this study, we observed several amino acid interactions between the P and S domains in the resting P domain conformation, while the P domain only connected *via* the extended flexible hinge in the rising P domain conformation. In addition, MNVs having a rising P domain showed a resolution about 2 Å lower (7.2 Å) than MNVs having a resting P domain (5.2 Å) (SI Appendix, Fig. S1*D* and *H*, SI Appendix, Fig. S4*F* and *J*). These facts suggest that MNVs with a rising P domain are relatively flexible. Such structural flexibility may facilitate viral assembly and disassembly. HNV GII.3 VLPs also showed that the rising P domain conformation easily collapsed due to surface tension of thin vitreous ice, thus leaving most of the particles with a resting P domain conformation in the thin ice of cryo-EM (SI Appendix, Fig. S14). These results suggest that the rising P domain conformation may represent an intermediate state of the viral assembly and disassembly in the host cells. Upon entering into the host cell, norovirus changes its structure to the metastable form to release the viral genome into cytosol at relatively lower metal ion concentration and higher pH conditions. Conversely, norovirus assembled in the host cell undergoes a conformational change to the stable form in a less amenable extracellular environment, such as the digestive tract, which is rich in free metal ions and with dramatically lower pH.

Recently, a dynamic feature of calicivirus capsid was reported in FCV. There, VP2, a minor capsid protein, forms a portal-like assembly on the capsid surface after receptor engagement, suggesting that it functions as a channel for delivery of the viral genome (18). In contrast to MNV, FCV requires a low pH for release of viral RNA from the capsid (34). It is unknown whether MNV has a similar portal structure consisting of VP2, but a rearrangement of the capsid structure, such as the dynamic rotation of the P domain, is believed to be a necessary step to form the portal and release the viral genome.

The interaction between P domain dimers has also been observed in human norovirus, where it is more complicated. The HNV GI.1 VLP shows dimer-dimer interactions at the P2 subdomain level, where the P2 subdomain of the C/C dimer links with four adjacent A/B dimers and the P domains rest on the S domains (7). By contrast, in HNV GII.10 VLP the similar dimer-dimer interactions are carried out at the P1 subdomain level because of the relatively small P2 subdomains, and the P domain rises from the S domain by ∼16 Å, like the rising P domain conformation in MNV. These different conformations of the P domains in different strains of HNV are only derived from the interaction sites at the P1 or P2 levels that can be structurally achieved by a simple rotation of the P domain. In addition, we previously reported a unique monoclonal antibody against the HNV GII.10 VLP, which binds to the occluded site between the P and S domains of the viral capsid and impairs binding to the histo-blood group antigens (13). This suggests that the antibody inhibits viral-cell attachment or/and infection by interfering with the rotation of the P domain from the rising position to the resting position. Currently, dynamic rotation of the P domain has been identified only in the infectious particles of MNV. If we are able to obtain infectious particles of HNV, we might be able to observe a similar phenomenon in human norovirus in the future.

## Materials and Methods

### Sample preparations of norovirus infectious particles and VLPs

Infectious particles of MNV-1 were produced using plasmid-based reverse genetics as previously described (21). Those of MNV-S7 were produced in the same procedure as MNV-1. pMuNoV-MNV-1 was constructed by exchanging the MNV sequence portion from MNV-1 to MNV-S7 with the In-Fusion cloning system (Takara Bio Inc.), according to the manufacturer’s protocol. MNV-1 infectious cDNA was kindly provided by I. Goodfellow as an MNV-1 cDNA plasmid (35). Infectious viruses of MNV-1 and MNV-S7 were propagated with RAW264.7 cells (American Type Culture Collection). The culture supernatant was collected two days after infection. MNV was precipitated using ultracentrifugation with a 30% sucrose cushion and purified with cesium chloride (CsCl) equilibrium ultracentrifugation. MNV was diluted with DMEM without any additional supplements and precipitated again to remove CsCl. The purified MNV pellet was resuspend in DMEM (Nacarai Tesque Inc.) or phosphate-buffered saline without calcium and magnesium containing 20mM EDTA at pH 8.0 (PBS(-)-EDTA (pH 8.0)) to prepare 10^7^ infectious virions /µL. VLPs of MNV-1, MNV-S7, and HNV GII.3 TCH04-577 strains were produced by a baculovirus expression system (24, 25) and purified with CsCl equilibrium ultracentrifugation in the same procedure as infectious particles.

### Analysis of MNV one-step propagation curve

RAW264.7 cells were cultured at a density of 10^6^ cells/well in 48-well plates and infected with the purified infectious particles of MNV-1 or MNV-S7 at a MOI = 2 or more. After 30 min of incubation at 37°C, the inoculum was removed, and the cells were washed three times with DMEM to remove unbound viruses, and DMEM containing 10% FBS (Thermo Fisher Scientific) was added. The plates were incubated at 37°C for the stated time; time zero indicates the time at which the medium was added. After 0, 3, 6, 9, 12, 15, 18 and 24 h of incubation, 20 µL of the culture medium was sampled and centrifuged at 10,000 g for 10 min at 4°C to remove cells and cell debris. 15 µL of supernatant was collected and stored at −80°C until RNA extraction. 10 µL of the supernatant was used for RNA extraction and qRT-PCR, according to published protocols (36). These assays were performed three times independently and calculated standard deviations (SD) plotted at each point.

### Evaluation of MNV-1 Binding using FACS

MNV-1 binding to the host cells was examined by FACS (fluorescence activated cell sorter) analysis. Briefly, 1 × 10^6^ of HEK293T/CD300lf cells were incubated with purified infectious MNV-1 particles (1×10^9^ CCID50), that were suspended in DMEM-10% FBS or PBS (-)-20mM EDTA (pH 8), for 30 min at 4°C. After washing, the cells were incubated with anti- MNV VP1 rabbit polyclonal antibody labeled by Dylight 488 (LNK221D488, BioRad) for 30 min at 4°C. The solution in each step includes 3% FCS and 20 mM NaN_3_ to prevent the cells from virus uptake. Afterward, the cells were washed again and analyzed by BD FACSMelody (BD Bioscience) and analyzed by using FlowJo software.

### Evaluation of cell binding and early genome replication of MNV in the host cells using qRT-PCR

RAW264.7 or HEK293T/CD300lf cells were cultured at a density of 10^5^ cells/well in 96-well plates and infected with the purified infectious particles of MNV-1 or MNV-S7 at a MOI ≦ 10. After 30 min of incubation at 37°C, the cells were washed twice with DMEM to remove unbound viruses. The total RNA was extracted by NucleoSpin RNA (Takara Bio Inc.) and the virus particles attached on the cell surface were estimated by qRT-PCR, according to published protocols (36). The early genome replication in each cell incubated for 0, 2, 4, 6, 8, 10, and 12 hours post-infection. These assays were performed three times independently and calculated standard deviations (SD) values and plotted at each point.

### Cryo-EM and image processing

Aliquots (2.5 µL) of the purified MNV infectious particles and HNV GII.3 VLP were placed onto R 1.2/1.3 Quantifoil grids (Quantifoil Micro Tools) coated with a thin carbon membrane that were glow-discharged using a plasma ion bombarder (PIB-10, Vacuum Device Inc.) immediately beforehand. These grids were then blotted and plunge-frozen using a Vitrobot Mark IV (FEI Company) with the setting of 95% humidity and 4°C. Vitreous ice sample grids were maintained at liquid-nitrogen temperature within a JEM2200FS electron microscope (JEOL Inc.), using a side-entry Gatan 626 cryo-specimen holder (Gatan Inc.), and were imaged using a field-emission gun operated at 200 kV and an in-column (Omega-type) energy filter operating in zero-energy-loss mode with a slit width of 20 eV. Images of the frozen hydrated norovirus particles were recorded on a direct-detector CMOS camera (DE20, Direct Electron, LP) at a nominal magnification of 40,000×, corresponding to 1.422Å per pixel on the specimen. Using a low-dose method, the total electron dose for the specimen is about 20 electrons per Å^2^ for a 3-second exposure. Individual images were subjected to per-frame drift correction by a manufacturer provided script.

For MNV-1 infectious particles suspended in DMEM and PBS(-)-EDTA (pH 8.0), 6,708 and 4,704 particles were selected from 1,046 and 2,188 images, respectively, and then extracted using RELION 2.0 (37) after determining the contrast transfer function (CTF) with CTFFIND4 (38). Alignment and classification of extracted particles were performed, and a 3D map was reconstructed in RELION 2.0 by using an initial model that generated with icosahedral symmetry by EMAN1 (39). The 3D reconstructions were computed, and the final resolutions of the density maps were estimated to be resolutions of 5.3 and 7.3 Å, respectively, using a gold standard Fourier shell correlation (cutoff 0.143) between two different independently generated reconstructions (40). 3D renders of the maps were created in UCSF Chimera (41). For MNV-S7 infectious particles suspended in DMEM and PBS(-)-EDTA (pH 8.0), 2,049 and 2,739 images were collected, respectively, and the 3D reconstructions were calculated with 17,820 and 5,063 particles at 5.2 and 7.2 Å resolution, respectively. For HNV GII.3 VLP containing the resting P domain and the rising P domain, 106 and 1,917 images were collected, and 3D reconstructions were calculated with 279 and 1,482 particles at 9.3 and 12.9 Å resolution, respectively. Data collection and image processing are summarized in Table S1.

For high-resolution structural analysis, cryo-EM images for MNV-S7 VLP were acquired with a Falcon II detector at a nominal magnification of 75,000, corresponding to 0.86 Å per pixel on a Titan Krios at 300 kV (Thermo Fisher Scientific). A low-dose method (exposures at 5 electrons per Å^2^ per second) was used, and the total number of electrons accumulated on the sample was ∼40 electrons per Å^2^ for an 8 second exposure. A GIF-quantum energy filter (Gatan Inc.) was used with a slit width of 20 eV to remove inelastically scattered electrons. Individual micrograph movies were subjected to per-frame drift correction by MotionCor2 (42). Particles were selected from the 2,746 images and the final 3D reconstruction was computed with 41,847 particles. The resolution of the density map was estimated to be 3.5 Å using the “gold standard” FSC criterion. Data collection and image processing are summarized in Table S2.

### Atomic model building of MNV-S7 VP1

The P domain atomic model of MNV-1 (PDB ID: 3LQE) and the S-domain atomic model of HNV (PDB ID: 1IHM) were used as templates for the homology model building of the P and S domains of the MNV-S7 VP1, respectively. Multiple-sequence alignments of VP1s of MNV-S7, MNV-1 and HNV were performed using the PROMALS3D program (43). The sequence alignments of the P and S domains were used as the input of MODELLER software (44) to generate the comparative models. The maps containing A, B, and C monomers were extracted using UCSF Chimera (41), respectively, for VP1s of MNV-S7 and MNV-1. The models were manually re-built in the individual maps from the homology model mentioned above using COOT (45) and refined using PHENIX (46). The models fitted to the cryo-EM maps were evaluated with the cross-correlation coefficient between the model and the cryo-EM map respectively, using “Fit in Map” of UCSF Chimera (41). Data collection, image processing, and model statistics are summarized in Table S2.

### Binding experiment of CD300lf to MNV-S7 VLP

Recombinant CD300lf (rCD300lf) (47) was kindly provided by S.Y. Park and prepared as reported (21). MNV-S7 VLP was incubated with rCD300lf for 1 h at room temperature, and then applied to a Quantifoil grid and plunge-frozen. Image data were collected and processed by the same procedure as the MNV virions. The final 3D map was reconstructed at 4.75Å resolution (gold standard FSC criterion) with 26,124 particles by imposing icosahedral (I1) symmetry. Then, the densities containing the periphery of the P2 subdomains of A/B and C/C dimers were extracted using UCSF Chimera (41), respectively. The extracted densities were used to generate masks around the P2 subdomains of A/B and C/C dimers using “Mask creation” command in RELION. Using the “relion_particle_symmetry_expand” command (37), the matrixes for each of the 60 subunits in each particle image were computed. The process generated 60 × 26,124 (1,567,440) sub-particle images. These sub-particles were subjected to “3D focused classification” with the masks around the P2 subdomains of A/B diner or C/C dimer, while keeping the orientations of each image used for the refinement of the 4.75-Å map. Among the six classes, one class (5.6% of the total) was selected that contained the density assumed to be bound CD300lf (SI Appendix, Fig. S13*A*). The particle images with the selected subunit were used for 3D refinement with a mask of the whole virus particle by imposing C1 symmetry. The map with the density of the CD300lf was reconstructed at 6.5Å resolution (gold standard FSC criterion) for total particle, and the resolution for only the focused area was estimated to be about 12Å (SI Appendix, Fig. S13*B*).

### *In silico* docking model of CD300lf to MNV-S7 capsid

The docking model of the CD300lf extracellular domain for the P domain dimer of MNV-S7 was constructed using Dock application of MOE 2015.10 (Chemical Computing Group). For the model of CD300lf, the crystal structure of the mouse CD300lf extracellular domain at a resolution of 2.1 Å (PDB ID: 1ZOX) was used. The program finally generated three potential docking models between the P domain and CD300lf, representing the docking scores based on generalized born solvation model of −52.39, −50.62, −50.61 kcal/mol, respectively. The model with the best score and physically acceptable poses was selected (SI Appendix, Fig. S13*D-F*).

## Data Availability

The data supporting the findings of this study is deposited to EMDB and PDB as described in Data Deposition section and can be obtained upon reasonable request from the corresponding authors.

## Acknowledgements

We thank T.J. Smith for providing the 8 Å cryo-EM map of MNV-1, T. Sato for helping the initial model building of the MNV-S7, K. Namba and Y. Kawaoka for their helpful discussions, and G.C. Howard and R.N. Burton-Smith for critical reading of the manuscript. We also thank S.Y. Park and M. Ohki for kindly giving us the recombinant CD300lf. This work was supported by AMED (to K.K.; Grant No. JP18fk0108034h and JP18am0101072j to Ka.M.), MEXT KAKENHI (Grant No. JP26102545 and JP16H00786 to Ka.M.), JSPS KAKENHI (to K.K.), and the collaborative programs for National Institute for Physiological Sciences (to K.K.).

## Footnotes

## Author Contributions

R.T., M.M., K.H., A.F., Ko.M., R.I., K.O., and K.K. prepared viral particles and performed cell adhesion experiments. C.S. and Ka.M. prepared cryo-EM specimens, performed 200 kV cryo-EM and image analysis, and built the atomic models. N.M. and K.I. collected and pre-processed data with the 300-kV microscope. M.Y. performed the docking simulation analysis. Ka.M. and K.K. supervised the work and coordinated experiments. C.S. and Ka.M. initially wrote the manuscript and prepared the figures with input from the other authors. All authors contributed to the finalization of the manuscript.

## Data deposition

Cryo-EM maps have been deposited in the Electron Microscopy Data Bank under accession numbers EMD-9735, EMD-9736, EMD-9737, EMD-9738, EMD-9739, EMD-9740 and EMD-9741. The atomic models have been deposited in the Protein Data Bank under accession number 6IUK.

