## Supplementary material for "Dynamic rotation of the protruding domain enhances the infectivity of norovirus": SI Appendix, Fig. S1 - S14

Kazuyoshi Murata

Kazuhiko Katayama

**This PDF file includes:**

Figures S1 to S14

Tables S1 to S2

Legends for Movies S1 to S2

**Other supplementary materials for this manuscript include the following:**

Movies S1 to S2

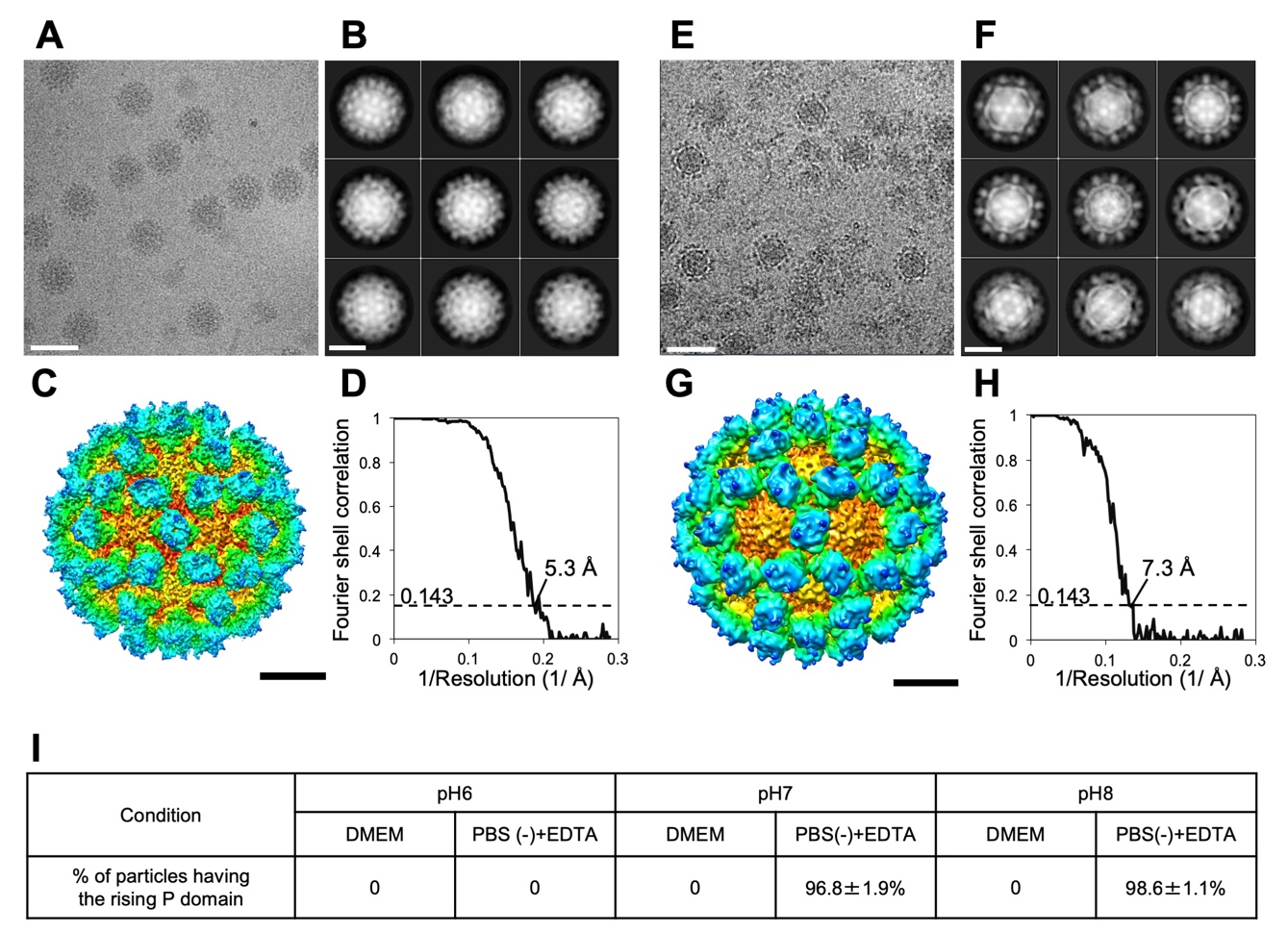

**Fig. S1.** Cryo-EM map generation of the MNV-1 infectious particles suspended in DMEM and PBS(-)-EDTA (pH 8). (*A* and *E*) Representative micrographs of MNV-1 in DMEM and PBS(-)-EDTA (pH 8), respectively. Scale bars, 500 Å. (*B* and *F*) The representative 2D class average images derived from *A* and *E*, respectively. Scale bars, 200 Å. (*C* and *G*) The surface-shaded depth-cued representations of the MNV-1 in DMEM and PBS(-)-EDTA (pH 8) viewed along the icosahedral twofold axis, respectively. Coloring is based on radii, as in Fig. 1*A*. Scale bars, 100 Å. (*D* and *H*) Plots of the gold-standard Fourier shell correlations (FSC) of the cryo-EM maps of MNV-1, respectively. Based on the 0.143 criterion for comparing two independent data sets, the image resolutions are estimated to be 5.3 and 7.3 Å, respectively. (*I*) The fraction of MNV-1 particles having the rising P domain conformation in different aqueous conditions. Random sampling of 1,000 particles was carried out five times under each condition.

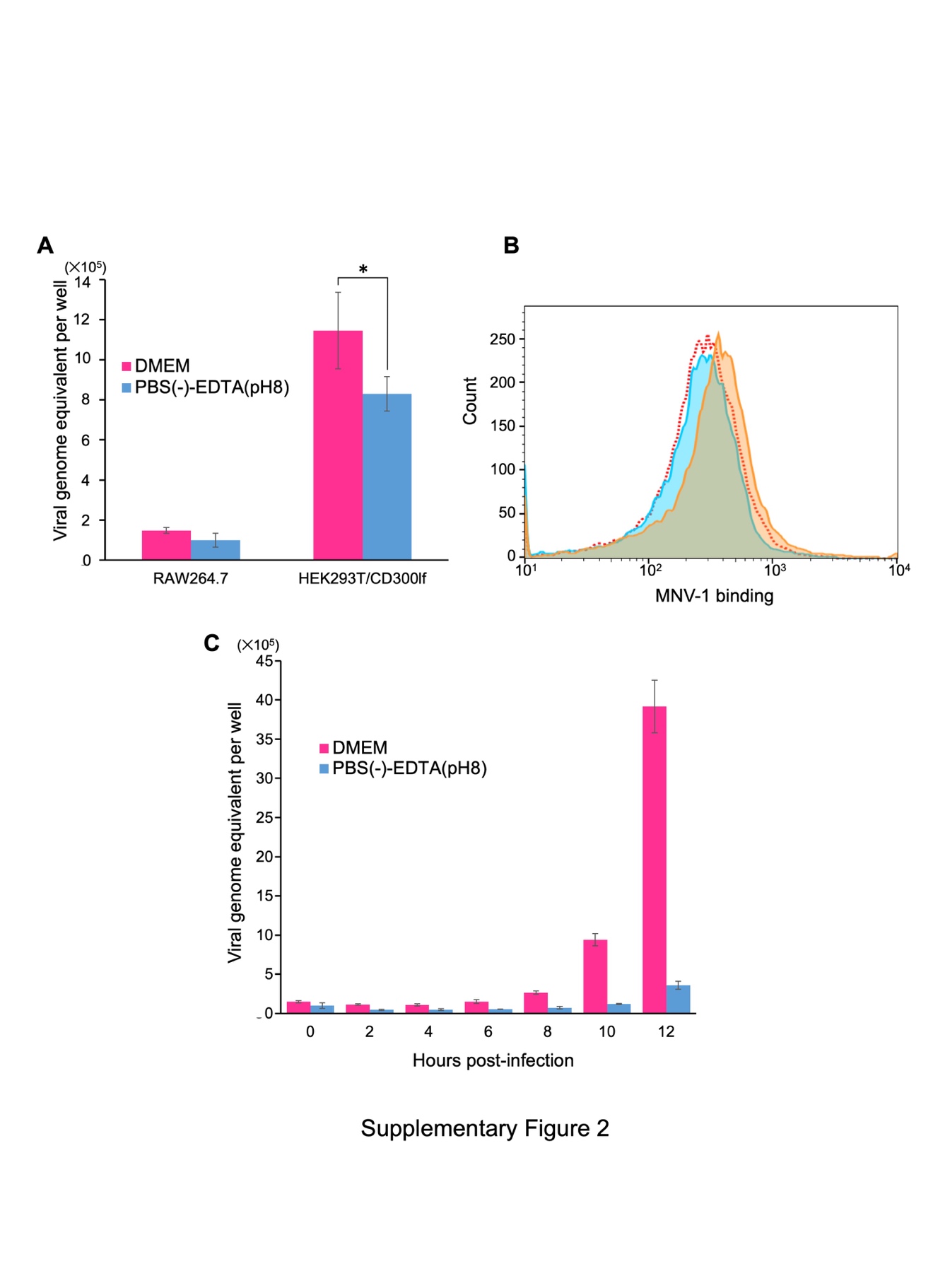

**Fig. S2.** Viral adhesion of MNV-1 particles to the host cells. (*A*) Viral initial attachment on the host cells at 30 minutes after post-infection of MNV-1 particles pretreated with DMEM and PBS(-)-EDTA (pH 8). After washing the cells with the solution, the viral RNA was extracted from viruses attached on the cells and the amount was estimated by qRT-PCR. *(p<0.05). (*B*) The cells were incubated with MNV-1 particles pretreated with DMEM (Orange), MNV-1 particles with PBS(-)-EDTA (pH 8) (Light blue), or without MNV-1 particles (Red dots). MOI was 10. After incubation, these cells were immunostained with anti-MNV VP1 antibody. X-axis represents the range of the MNV binding, and y-axis represents the number of the cells. Median and CV values are 357 and 275 (MNV-1 particles pretreated with DMEM), and 268 and 157 (MNV-1 particles pretreated with PBS(-)-EDTA (pH 8)), respectively. (*C*) Early genome replication of MNV-1 from 30 minutes to 12 hours. After removing the supernatant, the RAW264.7 cells infected with MNV-1 particles pretreated with DMEM or PBS(-)-EDTA (pH 8) were washed and collected at each time point of post-infection. The RNA was extracted and the amount was estimated by qRT-PCR.

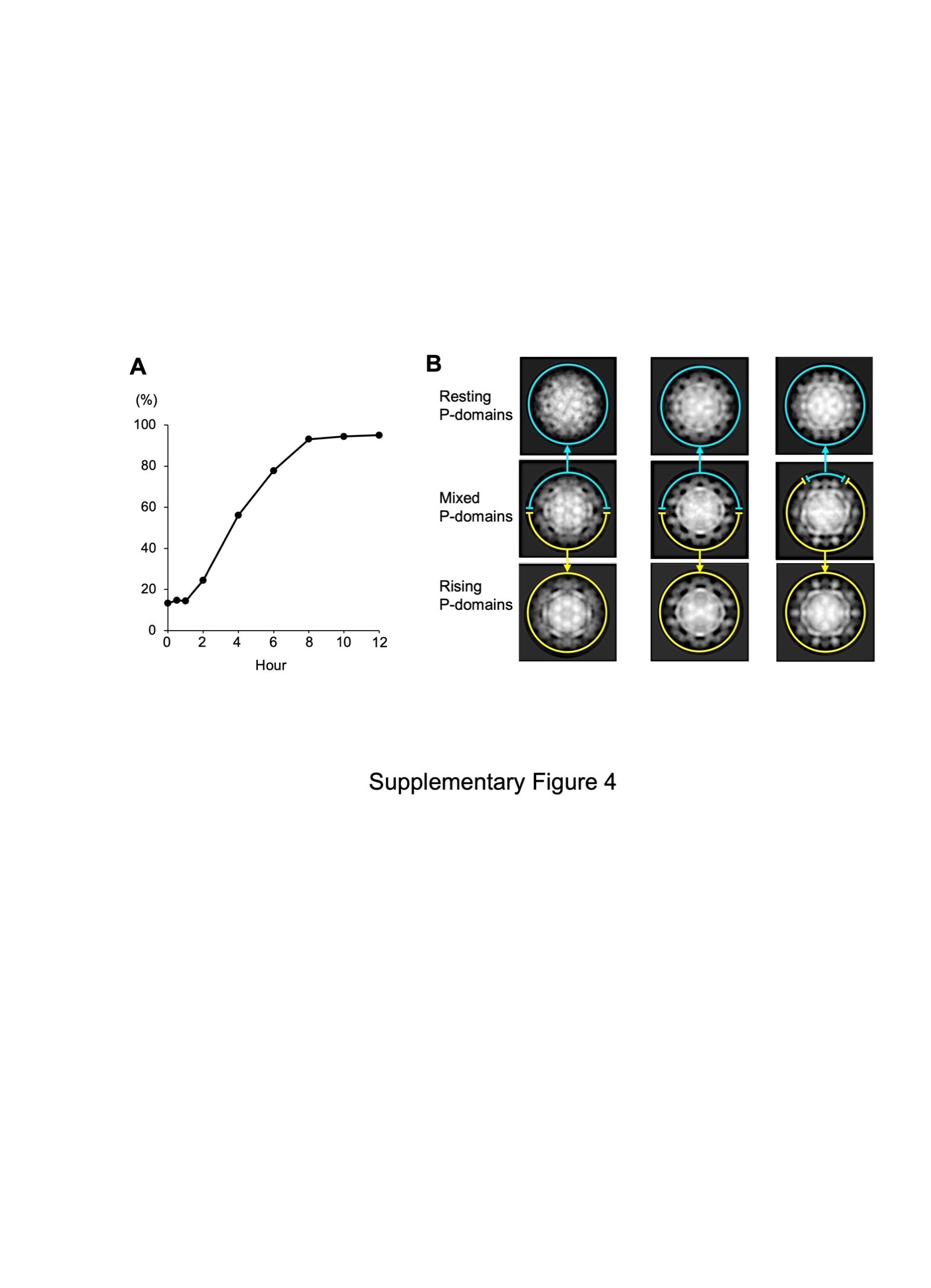

**Fig. S3.** Direct observation of the P domain’s conformational change in MNV-1 infectious particles. (*A*) MNV-1 particles having the rising P domain conformation in PBS(-)-EDTA (pH 8) were suspended in DMEM. The particles were then observed directly using cryo-EM. The percentage of the MNV-1 particles having the resting P domain conformation were plotted over time (n= ~200 particles in each point). (*B*) Representative 2D average images of MNV-1 particles between 2 and 6 hours. Particles mixed with two conformations were observed (second row panels). The mixed P domain structure within a single particle suggests that even if the conversion of individual P domains may be rapid, it would take time to modify the overall conformation of the capsid in this aqueous condition, where the P domains are connected together like a net.

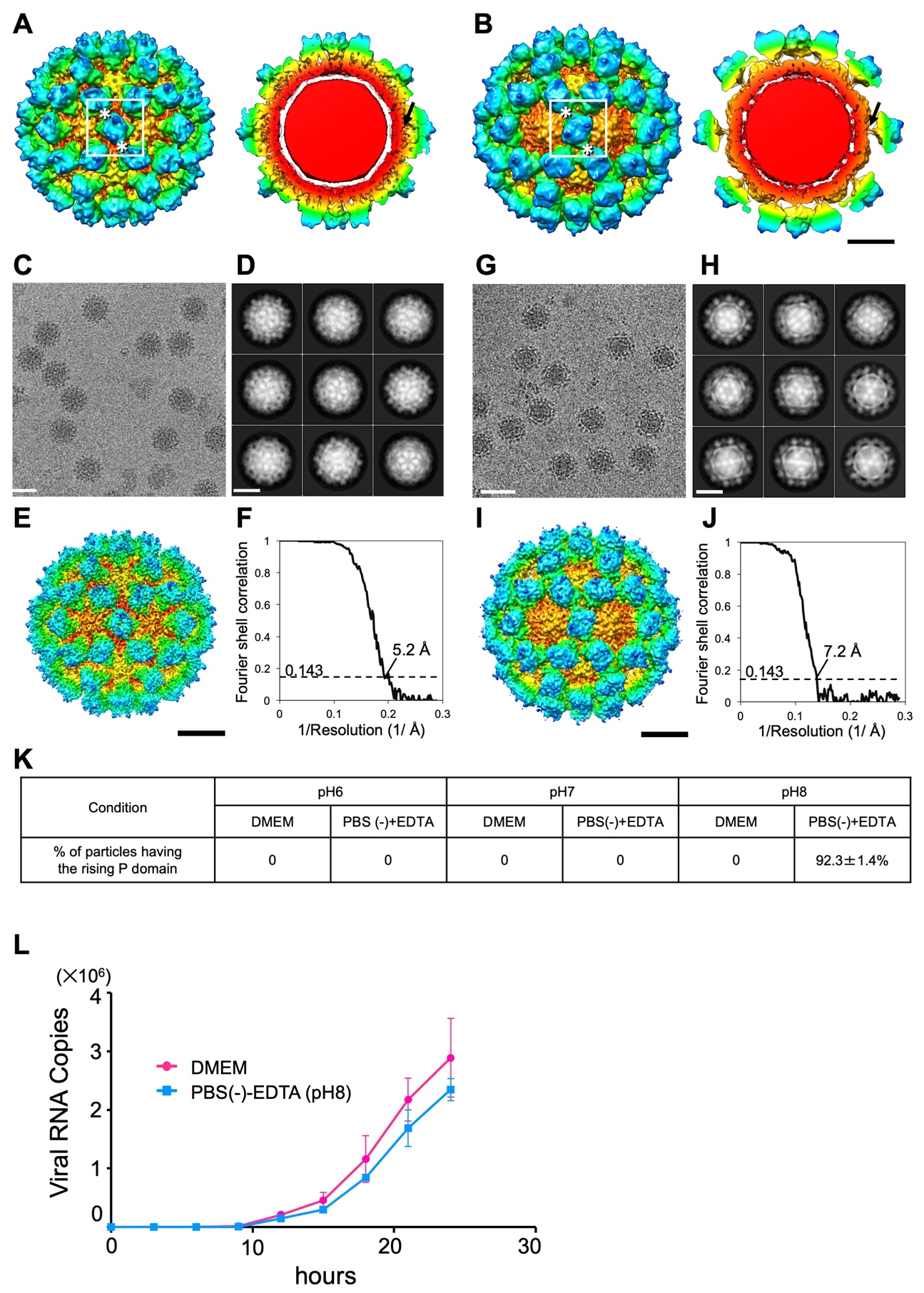

**Fig. S4.** Dynamic rotation of the P domains in the MNV-S7 infectious particle regulates the viral infection. (*A* and *B*) The cryo-EM structure of the MNV-S7 infectious particles, suspended in DMEM and PBS(-)-EDTA (pH 8), respectively, is low pass filtered to 8 Å resolution to highlight the P domain structure. Left and right panels show the isosurface display and center section, respectively. The P domain of MNV-S7 suspended in DMEM rests on the S domain (arrow in *A*) and interacts with the adjacent P domains at the outer P2 subdomain level (asterisks in *A*), called the resting P domain conformation. By contrast, the P domain of MNV-S7 suspended in PBS(-)-EDTA (pH 8) rises off the S domain (arrow in *B*) and interacts with the adjacent P domains at the inner P1 subdomain level (asterisks in *B*), called the rising P domain conformation. In the rising P domain conformation, all P domain dimers rotate ~70° clockwise, compared to the resting P domain conformation (white boxes), and the interaction sites are changed from the P2 to the P1 level. The P domain rising off from the S domain in PBS(-)-EDTA (pH 8) reversely rests on the S domain in DMEM. Scale bar, 100 Å. (*C* and *G*) Representative micrographs of MNV-S7 in DMEM and PBS(-)-EDTA (pH 8), respectively. Scale bars, 500 Å. (*D* and *H*) The representative 2D class average images derived from *C* and *G*. Scale bars, 200Å. (*E* and *I*) The surface-shaded depth-cued representations of MNV-S7 in DMEM and PBS(-)-EDTA (pH 8) viewed down the icosahedral twofold axis, respectively. Coloring is based on radii, as in Fig. 1*A*. Scale bar 100 Å. (*F* and *J*) Plots of the gold-standard Fourier shell correlation (FSC) of the cryo-EM maps of MNV-S7 (*E* and *I*) respectively. Based on the 0.143 criterion for comparing two independent data sets, the resolutions of the reconstruction are 5.2 and 7.3 Å, respectively. (*K*) The fraction of MNV-S7 particles exhibiting the rising P domain conformation in different aqueous conditions. Random sampling of 1,000 particles was carried out five times under each condition. The rising P domain conformation of MNV-S7 appeared at slightly higher pH than MNV-1 (Fig. S2*I*). (*L*) Propagation curves of MNV-S7 pretreated in DMEM and PBS(-)-EDTA (pH 8), respectively. The infectious particles were infected to RAW264.7 cells and measured the number of the duplicated viral RNA copies along the timeline by qRT-PCR. Error bars represent the standard deviations. MNV-S7 pretreated with PBS(-)-EDTA (pH 8) produced slower propagation of the virus than those pretreated with DMEM, though the difference was small compared to that of MNV-1 (Fig. 1*E*).

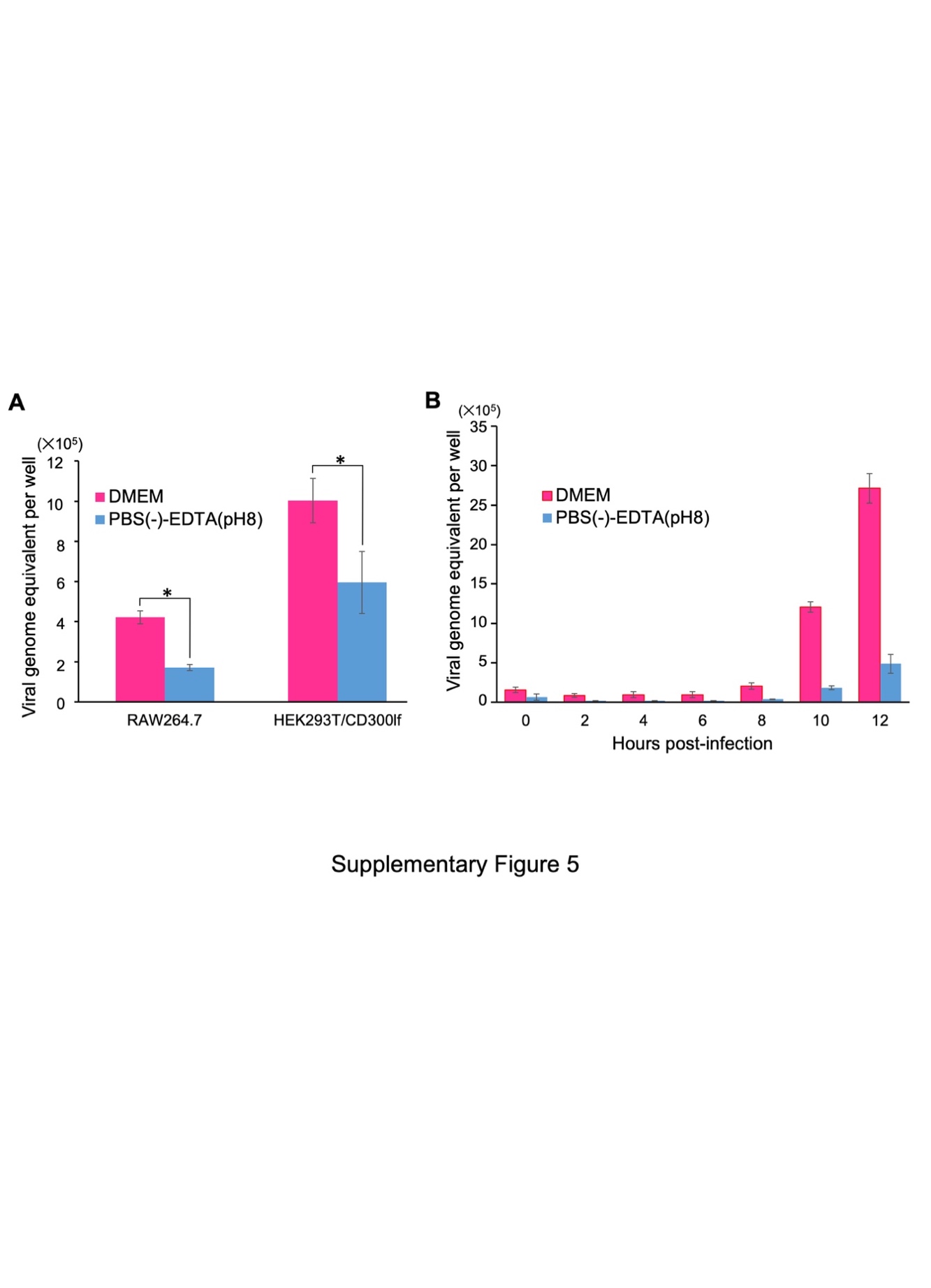

**Fig. S5.** Viral adhesion of MNV-S7 particles to the host cells. (*A*) Viral initial attachment on the host cells at 30 minutes after post-infection of MNV-S7 particles pretreated with DMEM and PBS(-)-EDTA (pH 8). After washing the cells with the solution, the viral RNA was extracted from viruses attached on the cells and the amount was estimated by qRT-PCR. *(p<0.05). (*B*) Early genome replication of MNV-S7 from 30 minutes to 12 hours. After removing the supernatant, the RAW264.7 cells infected with MNV-S7 particles pretreated with DMEM or PBS(-)-EDTA (pH 8) were washed and collected at each time point of post-infection. The RNA was extracted and the amount was estimated by qRT-PCR.

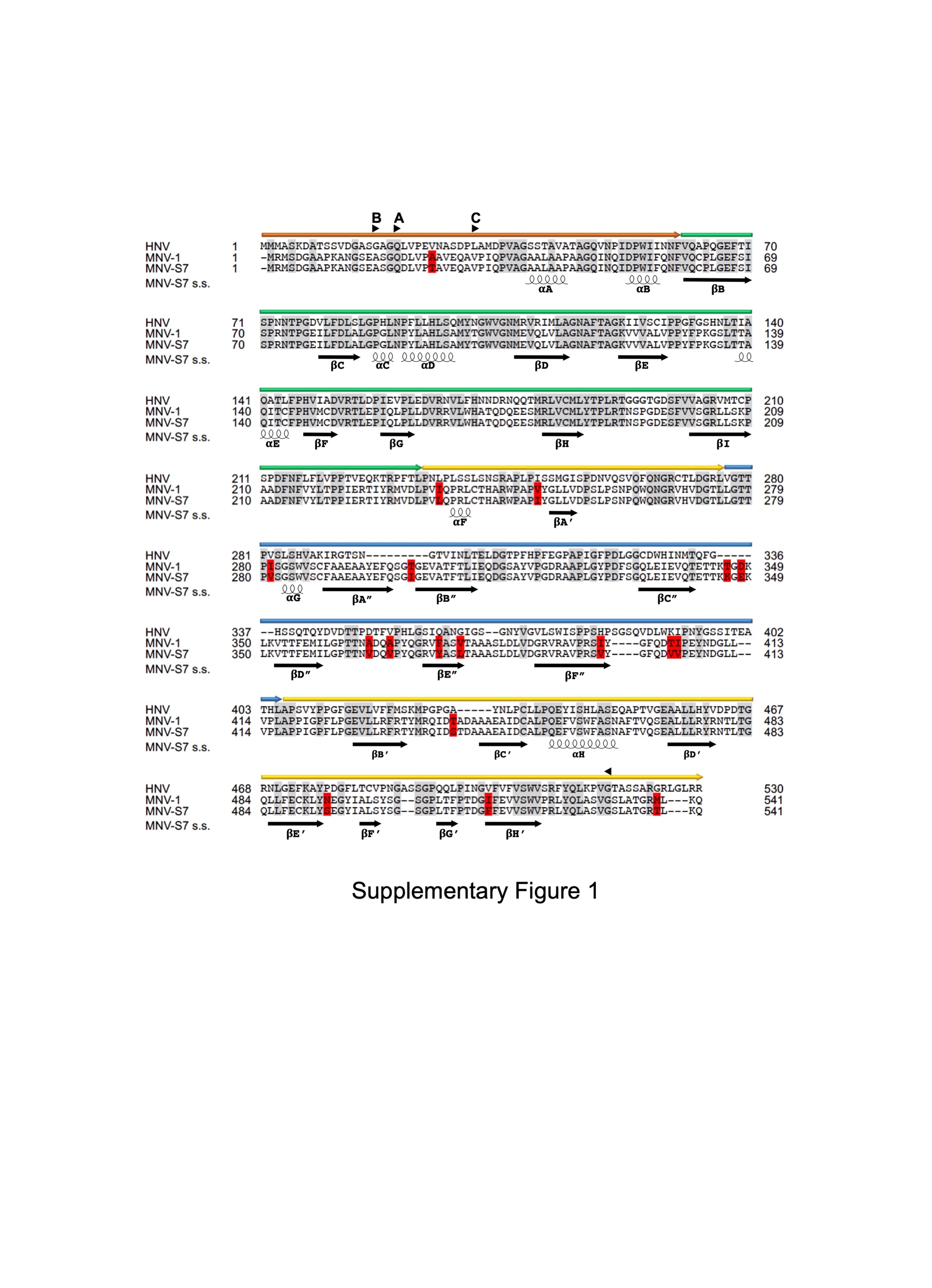

**Fig. S6.** Structural alignment of the VP1 protein in HNV(GI.1), MNV-1, and MNV-S7. For comparison, the amino acid sequences of VP1 of HNV (GI.1 Norwalk virus) and MNV-1 are aligned to that of MNV-S7. Crystallographic structures of the HNV S (PDB ID: 1IHM) and the MNV-1 P (3LQE) domains were used for the initial homology model building of MNV-S7. Orange, green, yellow, and blue arrows above the alignments represent the N-terminal, S domain, P1 subdomain, and P2 subdomain regions, respectively. Conserved sequences (46%) among HNV and MNV are indicated by gray, and different sequences (6%) between MNV-1 and MNv-S7 are indicated by red. Regions corresponding to major β-strands in norovirus capsid are represented by black arrows. Arrow heads indicate the start and end residues of the A, B, and C monomers to build the MNV-S7 structure model, respectively. MNV-S7 s.s. indicates the predicted secondary structures.

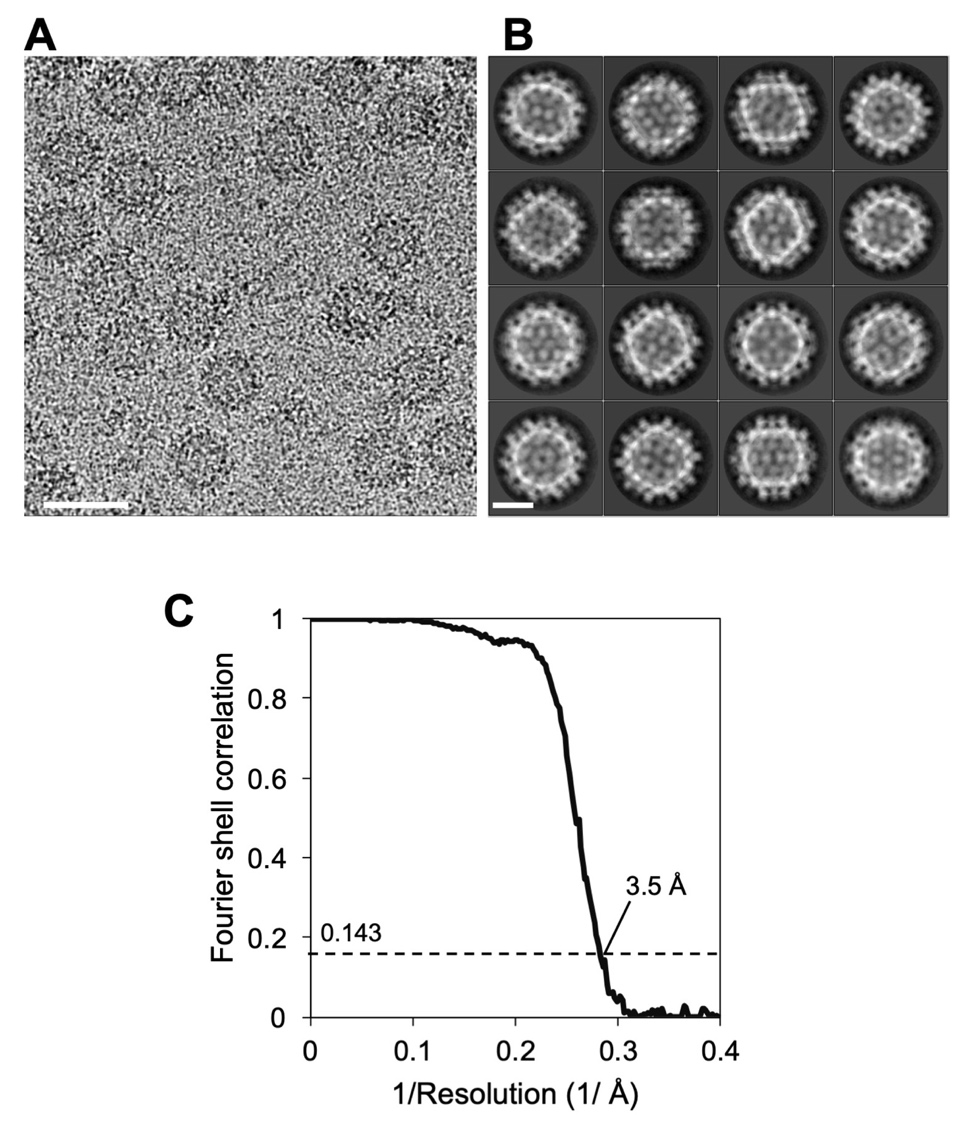

**Fig. S7.** High-resolution cryo-EM map generation of the MNV-S7 VLP suspended in DMEM by using a 300kV TEM. (*A*) Representative micrograph of the MNV-S7 VLPs. Scale bar, 500 Å. (*B*) An example of 2D class average images. Scale bar, 200 Å. (*C*) A plot of the gold-standard Fourier shell correlation. Based on the 0.143 criterion for comparing two independent data sets, the resolution of the reconstruction is estimated to be 3.5 Å.

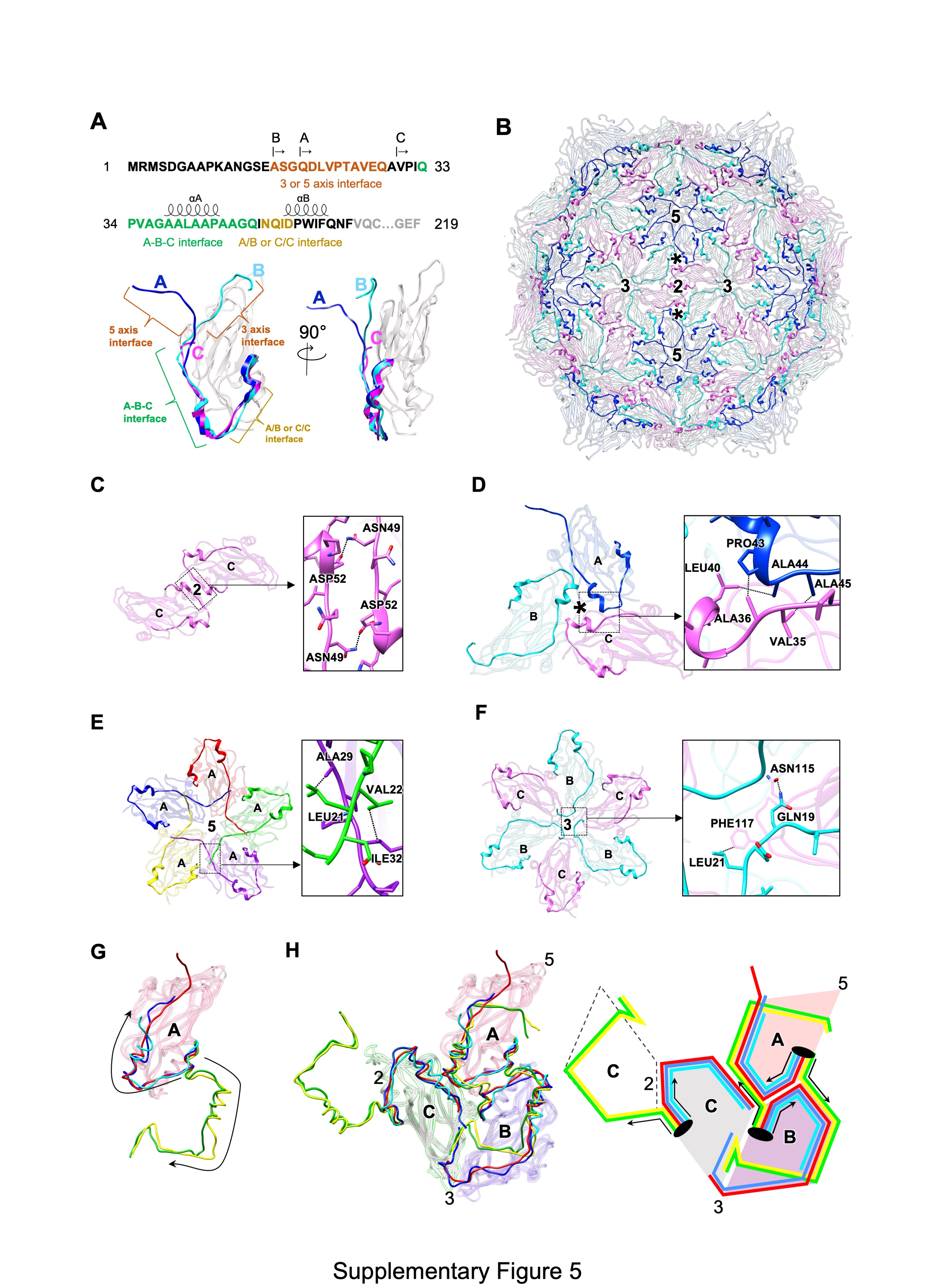

**Fig. S8.** Elaborate crosslinks between the S domains of VP1 in MNV-S7 VLPs. (*A*) Amino acid sequence corresponding to the N-terminal of the S-domain. The residues indicated by the arrows (Gln19, Ala16, and Val30) are the start residues of the sequences for modeling the respective VP1’s A, B and C monomers. Regions of the two short α-helices are indicated as αA and αB. In the lower panel, ribbon models of the S domain for the A, B and C monomers are superimposed. The residues colored by blue, cyan, and purple correspond to the N-terminals of the A, B, and C monomers, respectively. (*B*) N-terminals of the A, B and C monomers on the S domain capsid are highlighted with the same colors in *A*. Asterisks indicate the pseudo-threefold axis of the T=3 asymmetric units. (*C*) Interaction of the N-terminals of the S domains at the twofold axis. Asn49 and Asp52 of each C monomer form a charged interaction as shown by the dotted lines. (*D* and *E*) Interactions of the N-terminals at the pseudo-threefold axis in the T=3 asymmetric unit and the fivefold axis, respectively. The counter residues at the adjacent N-terminals interact by hydrophobic bonds as indicated by the dotted lines. (*F*) Interaction of the N-terminal at the threefold axis. The N-terminal edge of the B monomer interacts with the adjacent loop in the C monomer by charged and hydrophobic interactions as indicated by the dotted lines. (*G*) Comparison of the N-terminal arms (NTAs) of the S-domain in A monomer among the members of the *Caliciviridae* family: MNV (red), HNV GI.1 (blue, PDB ID: 1IHM), RHDV (cyan, PDB ID: 3J1P), SMSV (yellow, PDB ID: 2GH8) and FCV (Green, PDB ID: 3M8L). Arrows indicate the direction of the extended NTAs. (*H*) the adjacent B and C monomers were added in (*G*) to show the elaborate crosslinks between monomers. The right panel is a schematic figure of the left panel. The NTAs of S domain in MNV (red), HNV (blue) and RHDV (cyan) run along the outer bottom edge of their own S domain, while those in SMSV (yellow) and FCV (green) run along the outer bottom edge of their neighboring S domains.

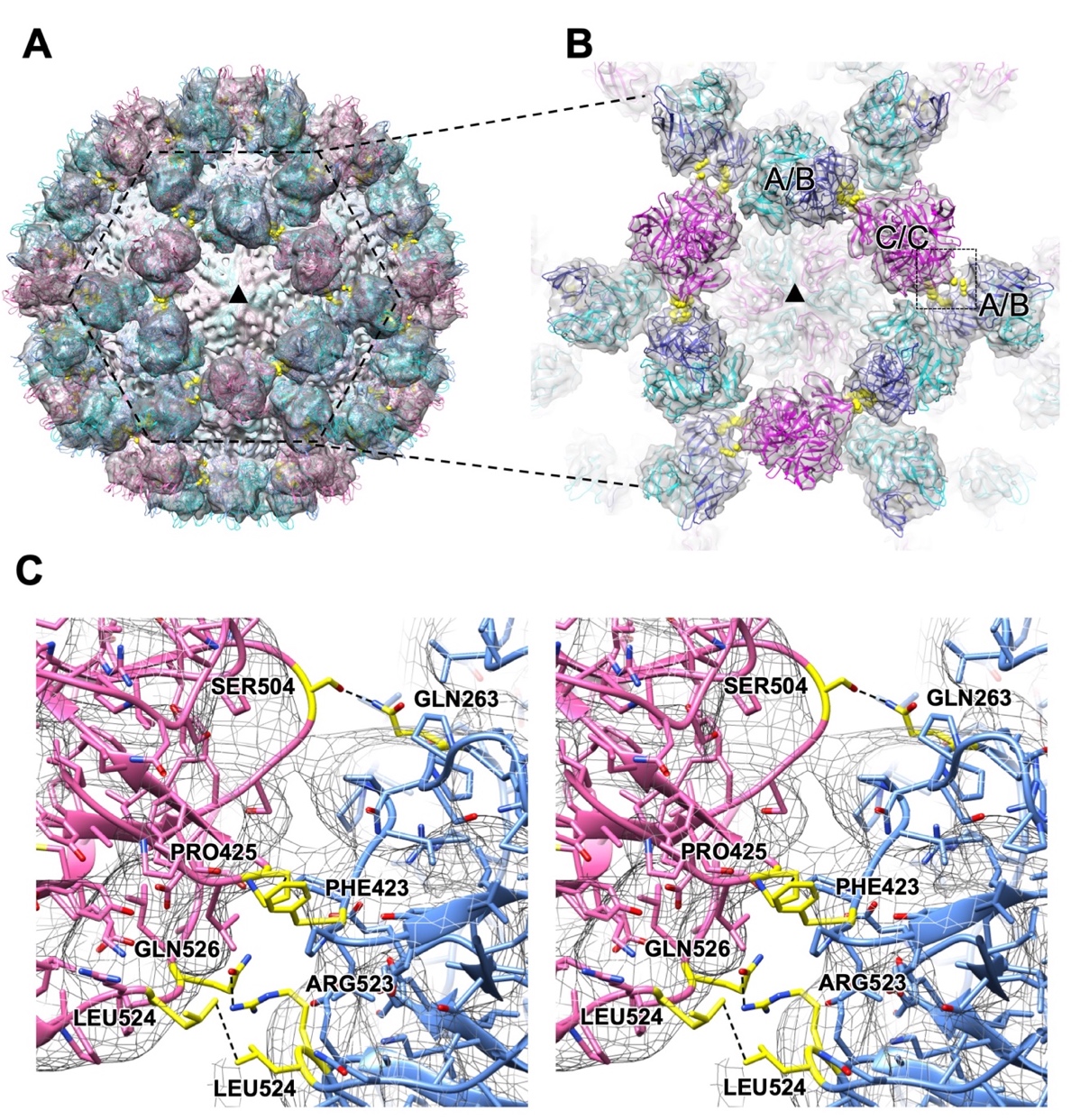

**Fig. S9.** Interactions between the rising P domain dimers. (*A*) The cryo-EM map of the MNV-S7 infectious particle suspended in PBS(-)-EDTA (pH 8) is shown on the threefold axis (black triangle). The residues forming interactions between the P domains are colored by yellow. C/C-dimers (purple) on the twofold axis form interactions with adjacent A/B dimers (blue) located around the fivefold axis. (*B*) The enlarged view of the dotted hexagon in *A*. (*C*) A stereo view of the candidate chemical interactions between the C/C and A/B dimers indicated by the dotted square in B. Pro425, Ser504, Leu524, and Gln526 of the C/C dimer are facing with Phe423, Gln263, Leu524, and Arg523 of the A/B dimer, respectively, where hydrophobic and charged interactions are suggested to be formed.

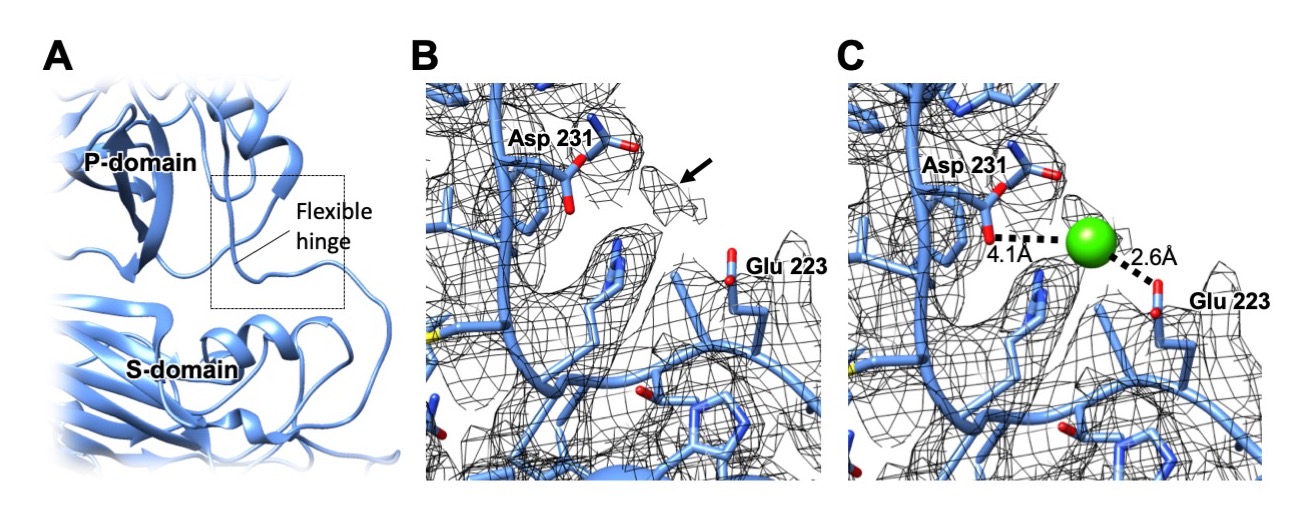

**Fig. S10.** The map and the molecular model of the flexible hinge in the resting P domain conformation. (*A*) An enlarged view of the flexible hinge connecting the S and P domains. The flexible hinge was indicated by the dotted box. (*B*) The enlarged map and model view of the box in *A*. Arrow indicates a density between Glu223 and Asp231. (*C*) Ca^2+^ as a possible metal ion is fitted into the density shown in *B*, and the protein structure around the metal is refined.

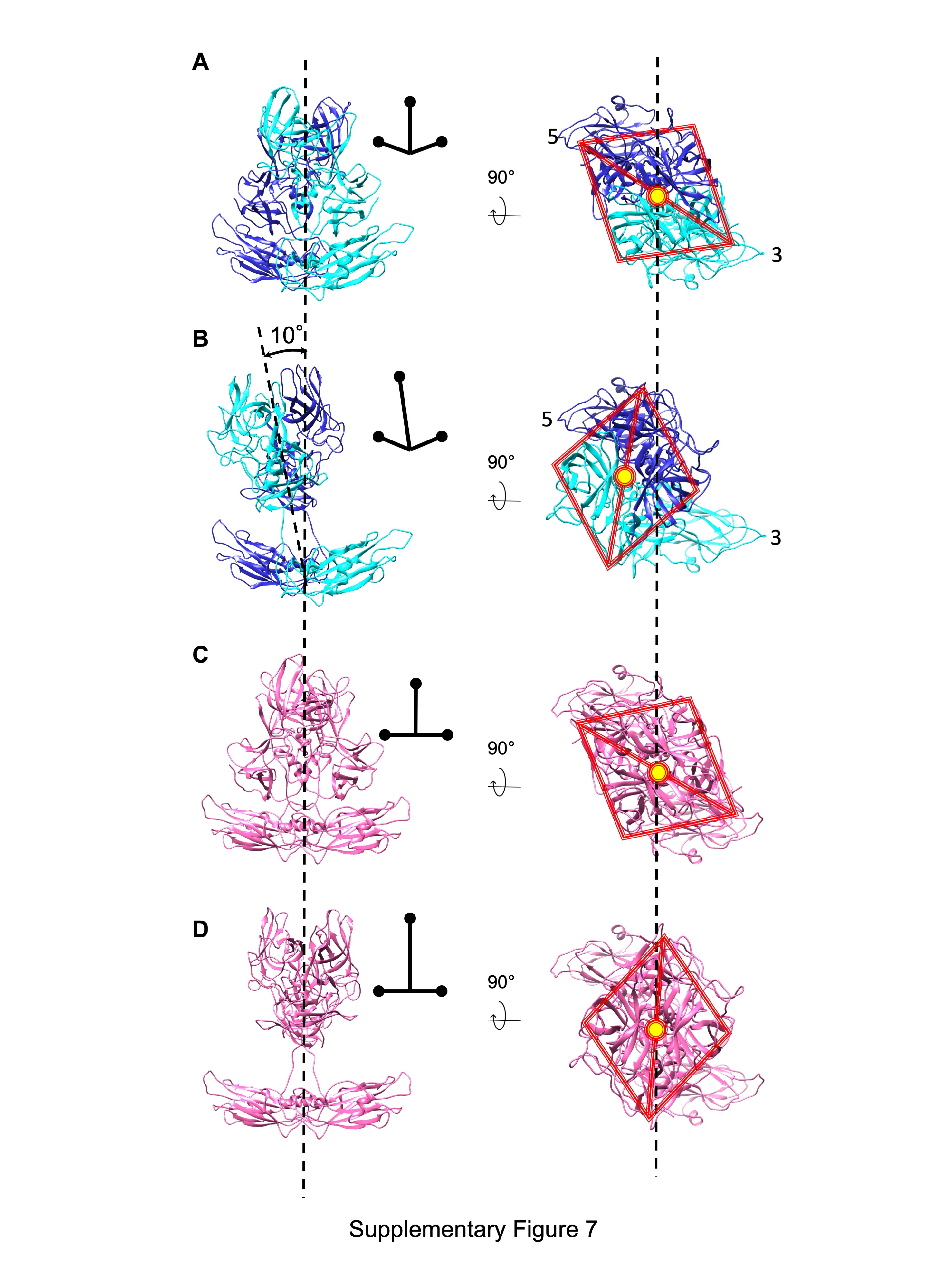

**Fig. S11.** The A/B and C/C dimers structures in the resting and rising P domain conformations. (*A and B*) The ribbon model of the A/B dimer in the resting and rising P domain conformation, respectively. The S domain dimer shows a bent conformation. The P domain dimer is tilted 10 degrees toward fivefold axis in the rising conformation. (*C and D*) The ribbon model of the C/C dimer in the resting and rising P domain conformations, respectively. The S domain dimer shows a flat conformation. The resting P domain dimer changes to the rising conformation without a tilt.

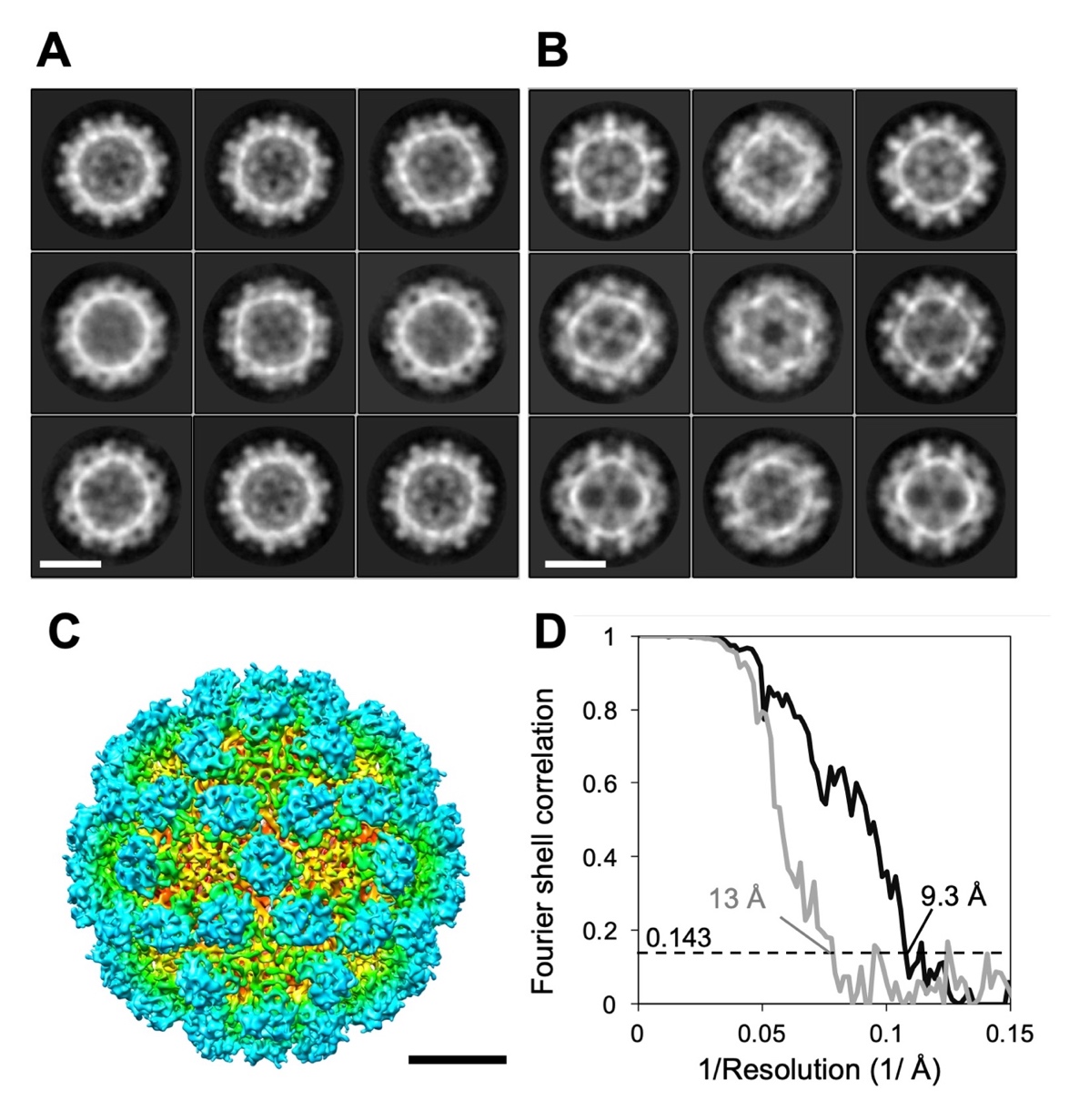

**Fig. S12.** Cryo-EM map generation of the HNV GII.3 VLP. (*A* and *B*) Representative 2D class average images of the HNV GII.3 VLPs with the resting and rising P domain conformations, respectively. Scale bars, 200 Å. (*C*) A surface-shaded depth-cued representation of the HNV GII.3 VLP with the resting P domain at 9.3 Å resolution. Viewed down the icosahedral twofold axis. Coloring is based on radii as in Fig. 1*A*. Scale bar, 100 Å. (*D*) Plots of the gold-standard Fourier shell correlation of the cryo-EM maps of the HNV GII.3 VLP with the resting and rising P domain conformations indicated by black and gray lines, respectively. Based on the 0.143 criterion for comparing two independent data sets, the resolutions of the reconstruction are 9.3 and 13 Å, respectively.

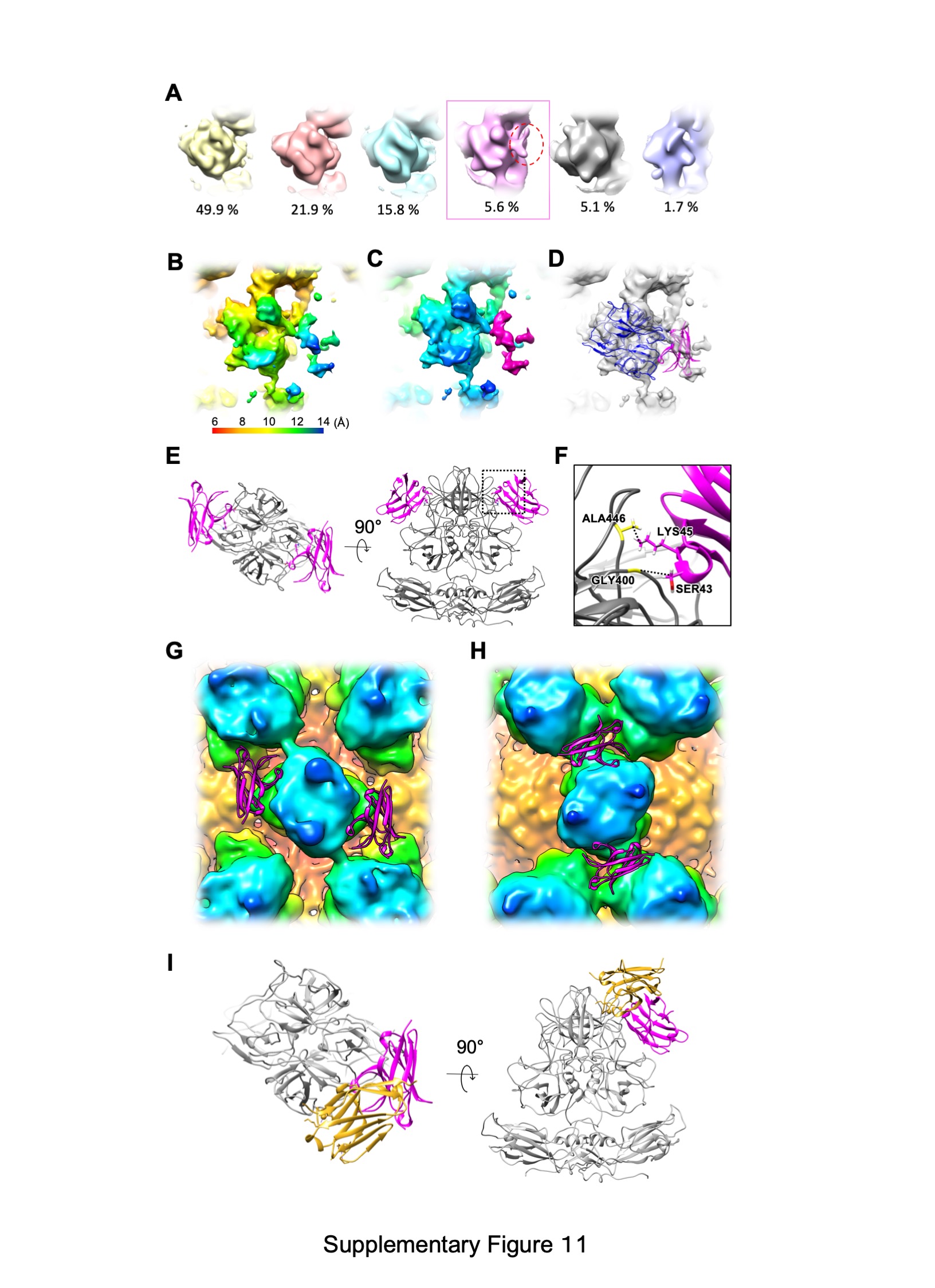

**Fig. S13.** Interaction of the P domain and a recombinant CD300lf (rCD300lf). (*A*) A focused classification result of the P-domain with rCD300lf. 1,567,440 sub-particle image capsid subunits generated from 26,124 particles were used. A class (5.6%) with a density assumed to be the receptor bound was selected for 3D refinement (red box). (*B*) Local resolution assessment in the 3D EM map of the P domain and rCD300lf. (*C*) Extra densities that appeared with the addition of rCD300lf are visualized by purple. (*D*) The Ribbon model of rCD300lf is fitted to the extra density using the docking simulation with a parameter set shown in Materials and Methods. (*E*) Protein-protein docking simulation between the P domain dimer (gray) and rCD300lf (magenta), based on the cryo-EM map in *D*. (*F*) Enlarged view of the square in *E* showing interactions of Gly400 and Ala446 in the βC'-βD' loop of the P domain (Fig. S4) and Ser43 and Lys45 in the C-C' loop in rCD300lf, respectively. (*G* and *H*) Possible binding modes of rCD300lf to the P domain in the resting and rising P domain conformations, respectively. A ribbon model of rCD300lf colored by magenta binds to the flat side of the P domain of MNV, based on the docking simulation. The binding site of rCD300lf is fully accessible in the resting P domain conformation (*G*), but it is restricted in the rising P domain conformation (*H*). An animation of rCD300lf accessing the P domain is shown in Movie S2. (*I*) A structural comparison with the X-ray crystallographic model (PDB ID: 5OR7) reported previously. In the co-crystal model, the actual binding site was close, thought rCD300lf approached from a higher position with a rotation of ~90°. The rCD300lf engaged a cleft between the A-B and D-E loops of the P2 subdomain in the crystal model (1), while the position in our model is slightly below the two loops. However, the spatial restriction of the CD300lf accessibility depending on the P domain rotation can be applied to the co-crystal model, although the binding modes are different.

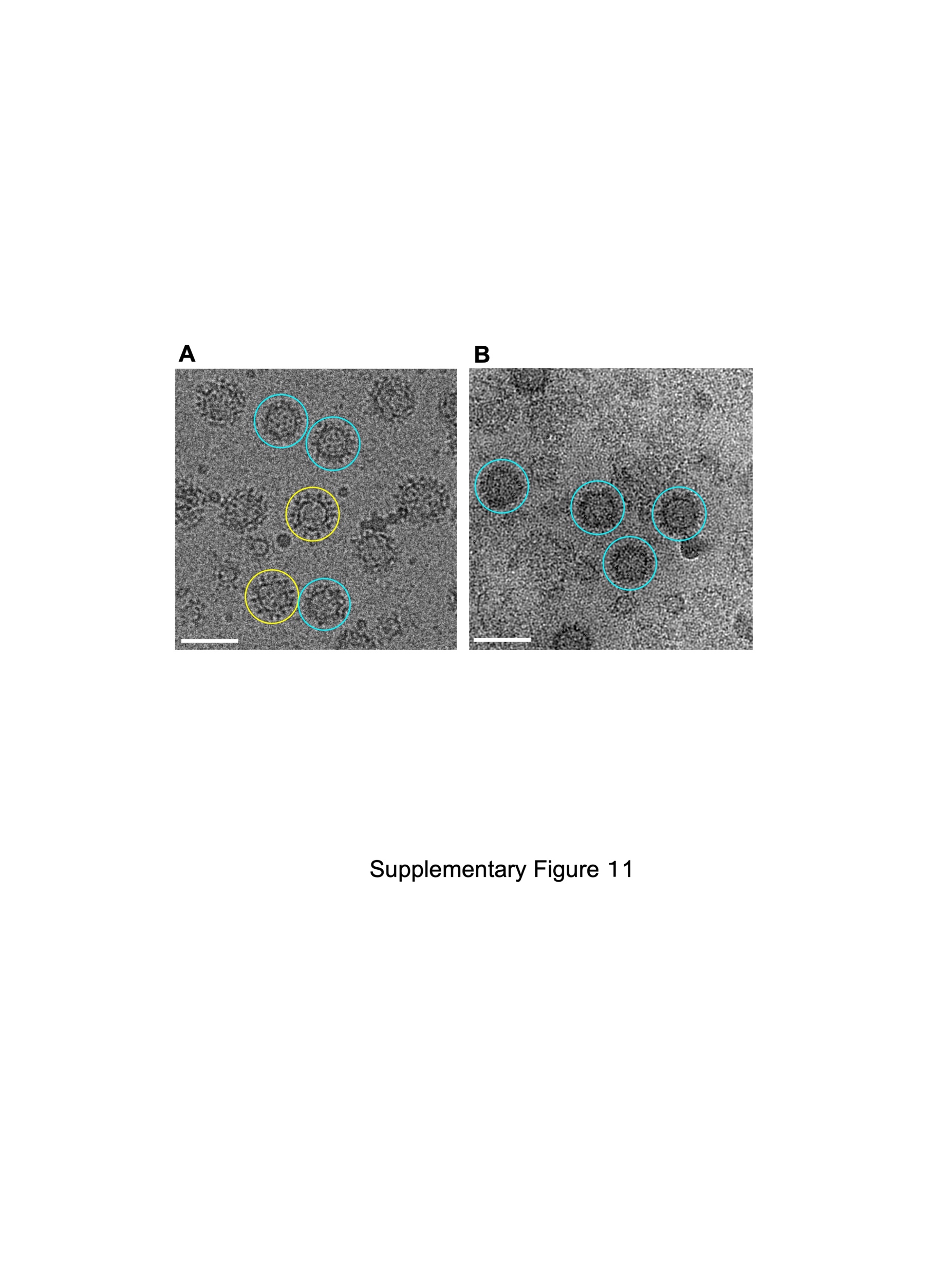
**Fig. S14.** Structural Stability of HNV GII.3 VLPs with the resting and rising P domain conformations. (*A*) Representative micrograph of the HNV GII.3 VLPs with the resting (blue circles) or rising (yellow circles) P domain conformations. (*B*) The VLPs having the rising P domain conformation easily collapsed due to the surface tension in thin vitreous ice, and only the stable VLPs with the resting P domain conformations were left.

**Table S1.** Data collection and image processing (MNV infectious particles, HNV GII.3 VLP, and MNV-S7 VLP with rCD300lf

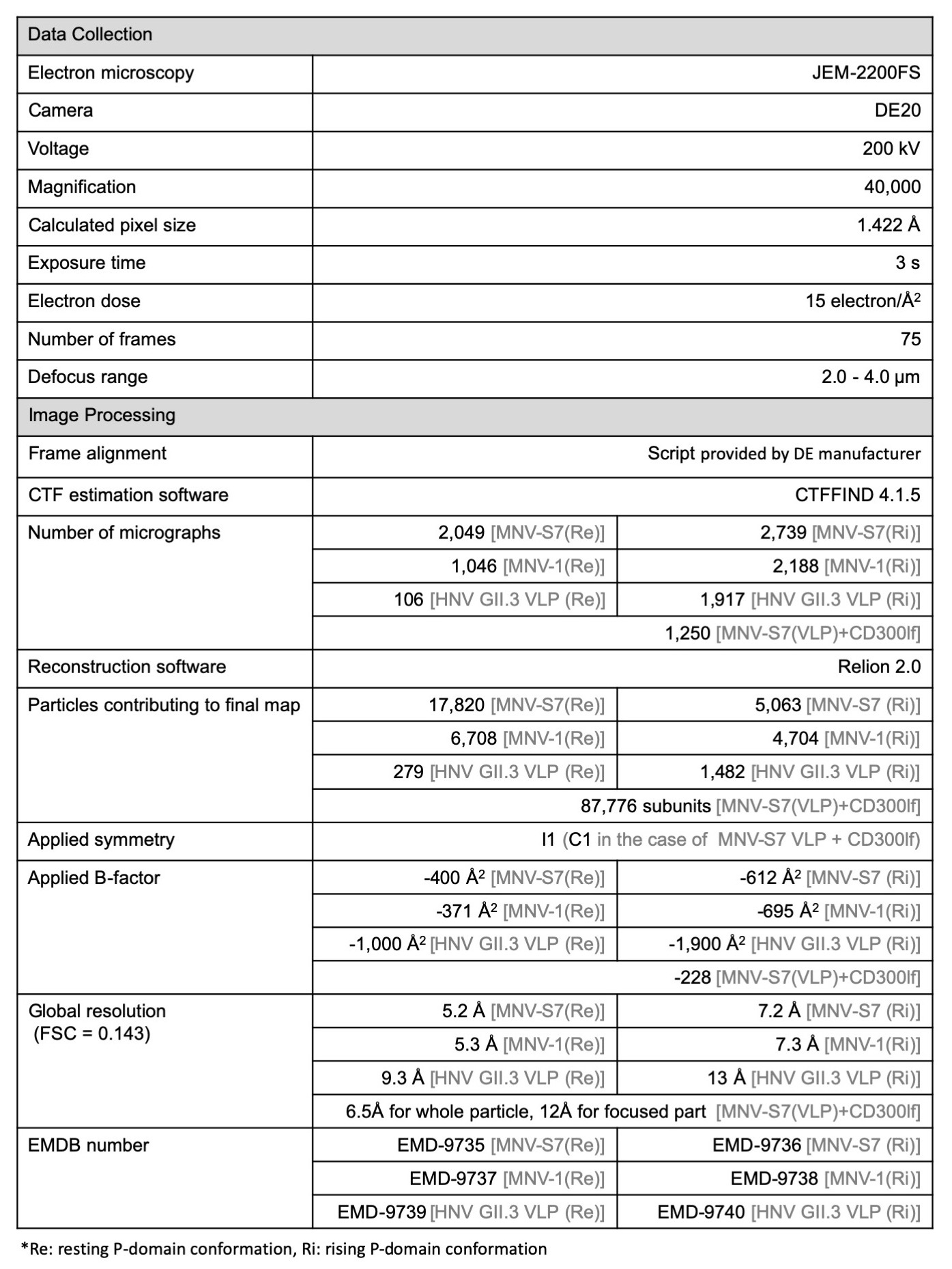

**Table S2.** Data collection, image processing and model statistics (MNV-S7 VLP)

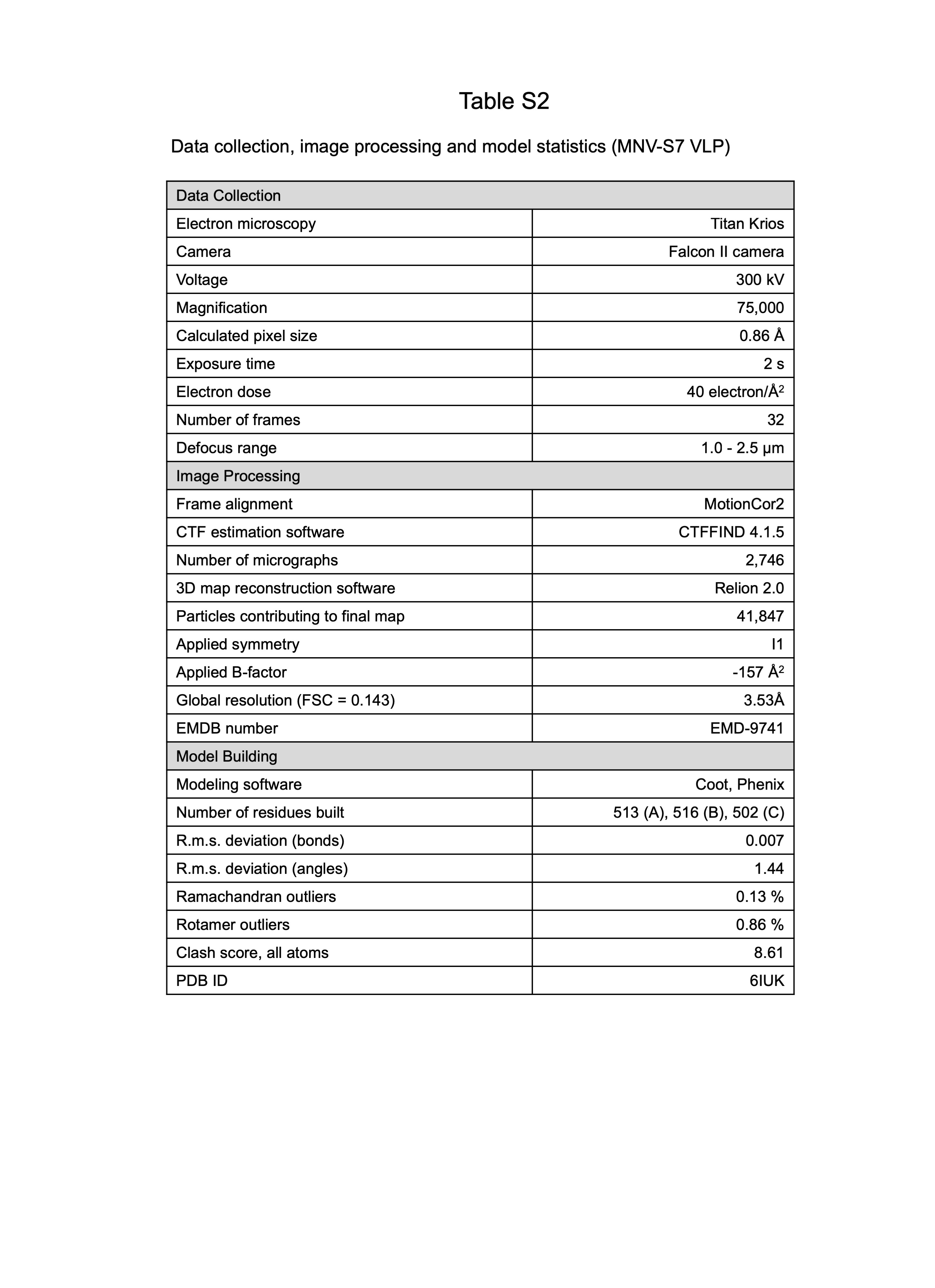

**Movie S1 (separate file).**

Molecular mechanism of the reversible rotation of the P domain dimers.

**Movie S2 (separate file).**

Accessibility of the cellular receptor molecule (rCD300lf) during the P-domain rotation.
